# Gut microbial-derived short chain fatty acids enhance kidney proximal tubule cell secretory function

**DOI:** 10.1101/2025.02.20.639317

**Authors:** Laura Giordano, Sabbir Ahmed, Thomas K. Van Der Made, Rosalinde Masereeuw, Silvia M. Mihăilă

## Abstract

The organic anion transporter-1 (OAT1), located at the basal side of kidney proximal tubule cells, is actively involved in metabolic waste excretion. During chronic kidney disease (CKD), the progressive decline in renal function, results in the accumulation of endogenous metabolites in the bloodstream, exacerbating comorbidities. CKD also leads to gut dysbiosis, resulting in an increased production of uremic metabolites and a reduced production of nephroprotective short chain fatty acid (SCFAs), mainly acetate, propionate and butyrate, thereby contributing to disease progression. Here, we studied the potential of SCFAs to enhance kidney function by modulating OAT1 activity, thereby facilitating uremic toxins secretion. Our findings demonstrate that propionate and butyrate significantly boost OAT1 activity by upregulating *SLC22A6/* OAT1 gene and protein expression, with butyrate enhancing the secretion of the uremic toxin indoxyl sulfate to the luminal compartment in our kidney-on-chip (KoC) system. Interestingly, SCFAs function independently of G-protein coupled receptor (GPCR) activation, inhibiting gene expression of class II histone deacetylases (HDACs). Transcriptome analysis suggests that such inhibition modulates cyclic AMP signaling pathway, activating *CREB1* and *PI3K* gene expression, both implicated in cell metabolism and resilience against stress, enhancing cellular fitness. These findings highlight the therapeutic potential of SCFAs in enhancing proximal tubule secretory activity, emphasizing their value as nutritional interventions in CKD management.

**HIGHLIGHTS:**

1. OAT1 activity is boosted by SCFAs via *SLC22A6* gene and protein upregulation
2. SCFAs act independently of GPCRs activation in proximal tubule cells
3. HDAC inhibition by SCFAs regulates cAMP pathway modulating *CREB1* and *PI3K* expression
4. SCFAs have the potential to serve as a therapeutic approach for kidney disease treatment

## 1. INTRODUCTION

The organic anion transporter-1 (OAT1) in kidney proximal tubule cells plays a pivotal role in eliminating metabolic waste, including endogenous toxins. This function is essential for maintaining kidney homeostasis. Dysregulation of OAT1 activity is increasingly recognized as a contributing factor in the progression of chronic kidney disease (CKD), highlighting the need for strategies to mitigate its dysfunction [1, 2]. Homeostasis is a dynamic and robust process tightly regulated by various factors to maintain physiological functions; consequently any dysfunction in one organ can directly affect another [3]. For instance, studies have highlighted the frequent appearance of gut dysbiosis among individuals suffering from kidney diseases compared to their healthy counterparts [4, 5] as a result of the bidirectional crosstalk between the gut and kidneys, named the gut-kidney axis [6]. Gut dysbiosis has been correlated with an increased production of microbiota-derived pro-inflammatory uremic toxins such as indoxyl sulfate (IS), para-cresol and trimethylamine N-oxide (TMAO) [5, 7]. As a result of kidney failure and an impaired excretion ability, these toxins start accumulating within the body [8] with detrimental effects on various organs and tissues.

The excretory mechanisms, mediated by the activity of membrane transporters located within the cell membranes of proximal tubule cells, can be classified into solute carrier (SLC) transporters or ATP-binding cassette (ABC) family members. The basolaterally expressed SLC transporter, organic anion transporter 1 and 3 (OAT1/3), encoded by *SLC22A6/8,* facilitate the uptake of organic anions; while their apical efflux is mediated by the ABC members, namely breast cancer resistance protein (BCRP/*ABCG2*) and the multidrug resistance-associated proteins 2 (MRP2/*ABCC2*) and 4 (MRP4/*ABCC4*), respectively. Previously, we demonstrated the ability of the kidneys to remotely detect and respond to increased plasma levels of uremic toxins via the modulation of transcription and expression of OAT1 involved in their excretion, also known as remote sensing hypothesis representative of the gut-kidney-axis [8].

Kidney failure substantially impact the bacterial genera in the intestine, as previously demonstrated both *in vivo* and clinically, with an increase in specific bacterial taxonomic groups upon the onset of CKD [4]. This genera shift is often accompanied by a depleted production of short chain fatty acids (SCFAs), mainly acetate, propionate and butyrate, which are microbiota-derived metabolites involved in safeguarding the intestinal barrier and exerting anti-inflammatory effects [5, 7]. Butyrate is predominantly absorbed in the intestine as a source of energy by colonocytes. Propionate and acetate are instead able to cross the portal vein, with propionate being substantially metabolized in the liver, while acetate is found within the systemic circulation, influencing several organs by regulating physiological functions [7]. SCFAs have been identified as being homeostasis modulators and exerting nephroprotective effects by modulating inflammation and immune responses, protecting from oxidative stress and ischemic injury [7]. Nonetheless, despite having been extensively studied in the intestine, limited studies have focussed on addressing their direct effect on human kidney cells, particularly on proximal tubule cells.

Accordingly, and as a result of their ability to mainly influence intestinal and energy homeostasis, immune and endocrine functions as well as epigenetic mechanisms [9], we hypothesized that acetate, propionate and butyrate might influence the activity of proximal tubule cells and could become a potential and safe therapeutic strategy to enhance renal tubular function resulting in improved elimination of metabolic waste. In this study, we investigated the impact of SCFAs on renal OAT1 functional activity. By employing a human conditionally immortalized proximal tubule epithelial cell line overexpressing OAT1 (ciPTEC-OAT1), we sought to provide insights into the potential regulatory mechanisms underlying SCFAs-OAT1 interactions and their implications for renal physiology and pharmacology. Understanding the interplay between SCFAs and OAT1 in renal epithelial cells may have significant implications for drug therapy and kidney health. Given the importance of OAT1 in renal drug handling and the physiological relevance of SCFAs, elucidating their interactions could lead to the development of novel therapeutic strategies targeting renal transport processes and metabolic regulation.

## 2. METHODS

### 2.1 Cell culture and exposure to SCFAs

The cell line ciPTEC-OAT1 was obtained from Cell4Pharma (www.cell4pharma.com) [10, 11]. Cells were cultured in Dulbecco’s Modified Eagle Medium/Nutrient Mixture (DMEM/F-12) without phenol red (Gibco, Life Technologies, Paisley, UK) supplemented with 10% (v/v) fetal calf serum (FCS) (v/v, Greiner Bio-One, Alphen aan den Rijn, The Netherlands), 5 μg/mL insulin, 5 μg/mL transferrin, 5 ng/mL selenium, 35 ng/mL hydrocortisone, 10 ng/mL epidermal growth factor and 40LJpg/mL tri-iodothyronine (Sigma Aldrich, Zwijndrecht, The Netherlands). Upon proliferation, cells were cultured at 33 °C with 5% (v/v) CO_2_ in T75 flasks up to 90% confluency and used between passage 50-60. For the experiments, cells were seeded at the appropriate density of 63,000 cell/cm^2^ and grown for 1 day at 33 °C, 5% (v/v) CO_2_ to allow for adherence and proliferation; and subsequently for 7 days at 37 °C, 5% (v/v) CO_2_ to allow for differentiation and maturation. Media was refreshed every other day. Upon complete maturation, cells were exposed individually or in combination to sodium-acetate (1-5 mM), sodium-propionate (1-5 mM) and sodium-butyrate (1-5 mM) (Sigma-Aldrich, Zwijndrecht, The Netherlands) for 24 h, referred to as acetate, propionate and butyrate, respectively.

### 2.2 Lactate-dehydrogenase cytotoxicity assay

The release of lactate-dehydrogenase (LDH) was measured using the Cytotoxicity Detection KitPLUS (Roche Diagnostics) following the manufacturer protocol. Upon 24 h exposure, supernatant was collected from each sample and incubated with 25 µL of the reaction mixture for 30 min at room temperature. Cells treated with Triton 1% were used as a positive control and untreated cells as negative control. Absorbance was measured at 490 nm using the GloMax™ Discover Microplate Reader (Promega, Leiden, The Netherlands). Arbitrary absorbance unit data were converted into percentage (%) and normalized to the positive control which was assigned as 100% cytotoxicity.

### 2.3 OAT1 activity fluorescein assay

To evaluate the effect of SCFAs exposure on OAT1 activity, an OAT-1 mediated fluorescein uptake assay was performed as previously described [12]. Upon exposure to SCFA at 1 mM either individually or in combination, ciPTEC-OAT1 were incubated with fluorescein (1 µM) prepared in HBSS for 10 min at 37°C. To confirm OAT1 activity, probenecid (500 µM, prepared in Krebs-Henseleit Buffer supplemented with 10 mM HEPES (KHH buffer, pH 7.4) a known inhibitor of OAT1, was simultaneously co-incubated with fluorescein for 10 min at 37°C. Uptake arrest was achieved by washing with ice-cold HBSS followed by cell lysis with 0.1 M NaOH. Intracellular fluorescence was quantified using the GloMax™ Discover Microplate Reader (Promega, Leiden, The Netherlands) at excitation and emission wavelength of 490 nm and 518 nm, respectively. Arbitrary fluorescence unit (AFU) data were normalized to the protein amount of each well, which was quantified using the Pierce BCA Protein Assay kit (Thermo Fisher). Protein amount was quantified using the GloMax™ Discover Microplate Reader (Promega, Leiden, The Netherlands) at absorbance 560 nm. Background values were subtracted from the arbitrary absorbance unit. Following normalization of the AFU to protein amount, data were converted into percentage (%) and the fluorescein uptake of untreated cells was used as positive control, reflecting 100% uptake.

### 2.4 Quantitative Real Time PCR

Total mRNA from ciPTEC-OAT1 seeded in 12 well-plates and exposed to acetate (1 mM), propionate (1 mM) and butyrate (1 mM) either alone or in combination for 24 h, was isolated using the RNeasy mini kit (QIAGEN, Hilden, Germany) according to manufacturer’s procedure. Following mRNA isolation, the mRNA concentrations and purity were assessed using the NanoDrop® ND-1000 spectrophotometer (Thermo Fisher Scientific). Reverse transcription of RNA to complementary DNA (cDNA) was performed using the iScript^TM^ Reverse Transcription Supermix (Bio-Rad Laboratories, Hercules, CA, USA) according to the manufacturer’s instructions. qRT-PCR (Real Time-quantitative Polymerase Chain Reaction) was subsequently performed to measure the relative gene expression of GPR41/*FFAR3*, GPR43/*FFAR2*, GPR109A/*HCAR2* and OAT1/ *SLC22A6* using SsoAdvanced Universal SYBR® Green Supermix (Bio-Rad Laboratories) and the CFX96TM Real-Time PCR Detection System (Bio-Rad Laboratories). Data were analyzed with the Bio-Rad CFX Manager^TM^ Software version 3.1 (Bio-Rad Laboratories) and displayed as relative gene expression, using untreated cells as the reference sample. ATP synthase (*ATPs*) was used as a housekeeping gene for normalization. Primer sequences are summarized in Table 1. Relative gene expression to control was calculated as fold changes using the 2−ΔΔCt method. All reactions were carried out with the equal amount of 60 ng/uL of cDNA sample.

**Table 1:**
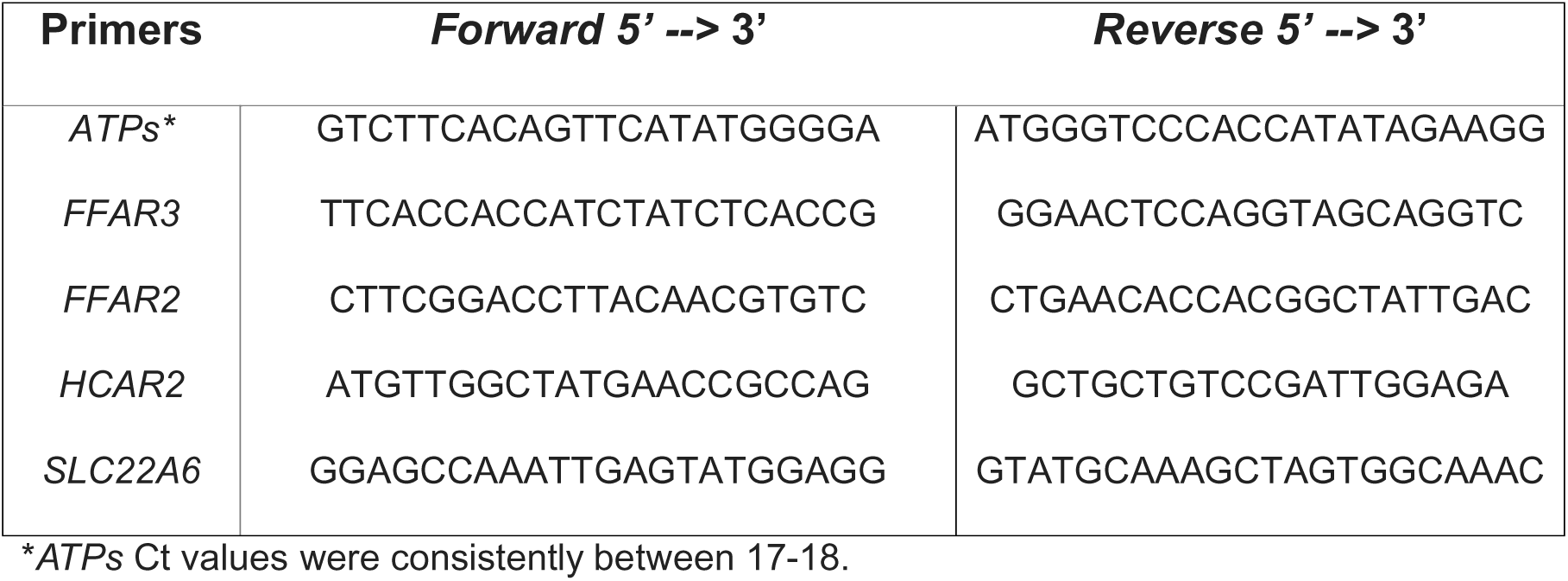
List of primers for qRT PCR analysis.

### 2.5 Western blotting

For protein expression analysis cells were washed once in ice-cold HBSS and subsequently lysed in ice-cold RIPA Lysis Buffer containing proteinase (Thermo Scientific, Vienna, Austria) for 30 min and centrifuged at 4°C. The obtained samples were then quantified using the BCA Protein Assay Kit (Thermo Scientific, Vantaa, Finland) and denaturated at 95°C for 5 min. From each sample, 20 µg were loaded and separated on a 4-20% polyacrylamide gradient SDS gels (Bio-Rad Laboratories, Hercules, CA), and transferred to a PVDF membrane (Bio-Rad Laboratories, Hercules, CA). The membranes were blocked using 5% milk in 0.1% Tween PBS (PBS-t) for 2 h and incubated overnight at 4°C with the primary antibody GPR41 (1:1000, Invitrogen, PA-99629), GPR43 (1:1000, Invitrogen, PA5-100944), GPR109A (1:1000, Invitrogen, PA5-90579) and OAT1 (1:1000, Invitrogen, PA5-26244) diluted in blocking buffer. β-tubulin (1:3000, Invitrogen, PA5-21826) was used as a housekeeping protein control. Thorough washing of the membranes was performed three times prior to incubation with the secondary antibodies anti-rabbit (1:5000, Dako, P0448, United States) performed in blocking buffer at room temperature for 1h. Membranes were subsequently washed three times and incubated for 5 min at room temperature in the dark with the ECL Prime Western Blotting Detection Reagent (Cytiva Amersham TM). Imaging was performed using the ChemiDoc™ MP Imaging System (Bio-Rad Laboratories, Hercules, CA) for band detection which was then quantified using the ImageJ software (version 1.53c, National Institutes of Health, United States) and normalized to the β-tubulin housekeeping protein expression.

### 2.6 Kidney-on-chip cell culture

The KoC culture was based on a previously established platform, which was slightly modified [13, 14]. 3D polylactide chambers were used. The inner (basal) part of the HFM represents the peritubular circulation compartment, whereas the outer (apical) part is representative of the glomerular filtrate compartment (Fig. 2A). Briefly, microPES hollow-fiber membranes (HFM) (inner diameter of 300 µm, outer diameter of 500 µm and pore size of 0.5 µm, 3M GmbH, Wuppertal, Germany) were cut and guided through the chamber using a steel wire. The bottom of the chamber was sealed with a 24LJ×LJ60LJmm glass cover slip (Menzel-Gläser, Braunschweig, Germany) using Loctite EA M-31CL glue, to let dry for 24 h and the interconnection between the sides reservoirs was then made leak tight with the biocompatible GI-MASK Automix glue (Coltene, Lezennes, France). Once ready, chambers were sterilized with UV light overnight and with 70% (v/v) EtOH for 30 min. Subsequently the HFM were coated with a double biomimetic coating of L-3,4-di-hydroxy-phenylalanine (L-Dopa, 2 mg/mL in 10 mM Tris buffer, pH 8.5) for 5h at 37LJ°C and 5% CO_2_, followed by human collagen IV (25LJµgLJmL^−1^ in PBS) for 2h at 37LJ°C and 5% CO_2_. In between each step, the chamber was washed 3× using sterile HBSS. CiPTEC-OAT1 were seeded in the chambers at a seeding density of 1LJ×LJ10^6^ ciPTEC-OAT1/HFM and were cultured for 3 days at 33LJ°C with 5% CO_2_ to allow for proliferation, followed by 7 days at 37LJ°C with 5% CO2 to allow for maturation. Culture medium was refreshed every 2–3 days.

### 2.7 FITC-inulin leakage assay and IS clearance using the KoC system

Once the bioengineered kidney tubules were experiment ready, they were placed on a 2D plate rocker at a speed of 1 rotation per minute and a 10° angle (VWR, Breda, The Netherlands) while being basally exposed to acetate (1 mM), propionate (1 mM) or butyrate (1 mM), either alone or in combination for 24 h. Following incubation, cells were washed once and subsequently exposed basally to IS (100 µM) and fluorescein isothiocyanate-inulin (FITC-inulin, Sigma Aldrich, Zwijndrecht, the Netherlands) (0.1 mg/mL) in serum free DMEM/F-12 for 4 h at the same speed and angle as for the 24 h incubation. FITC-inulin co-incubation with IS was performed to quantify for paracellular permeability and degree of monolayer formation. Unseeded HFM were used as a positive control for leakage. Supernatants from both the apical and basal compartments were collected and fluorescence measured at excitation wavelength of 492 nm and emission wavelength of 518 nm using the GloMax™ Discover Microplate Reader (Promega, Leiden, The Netherlands). Results are shown as permeability coefficient (P_f_):

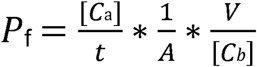

where: *[C_a_]* = apical concentration of FITC-inulin (mg/mL); *t* = the perfusion time (hours); *A* = the outer area of the HFM (cm^2^); *V* = the volume of the apical chamber (mL); *[C_b_]* = the basolateral concentration of FITC-inulin (mg/mL).

IS quantification was performed via LC-MS/MS by processing the same supernatants using a method previously developed in our lab [15]. Apical and basal compartments were washed once in ice cold HBSS whereas the inner part of the HFM was washed via perfusion with the help of a needle and syringe filled with ice cold HBSS. The HFM was then removed from the chip and placed in 0.1 M NaOH to lyse the cells. Protein quantification was performed using the Pierce BCA Protein Assay kit (Thermo Fisher) as described above.

### 2.8 RNA sequencing analysis

Cells were lysed in ice-cold RTL buffer (Cat. No. 79216, QIAGEN) and stored at −80°C prior to processing. RNA extraction was performed on a QIAsymphony isolation robot via QIAsymphony RNA Kit (931636, QIAGEN) and miRNA CT 400 protocol. The quality of the RNA was assessed with the RNA Kit (15 nt) (Cat. DNF-471-1000, Agilent, Santa Clara, CA, USA) using the Agilent Fragment Analyzer 5300 system, whereas RNA quantity was assessed with the Qubit RNA HS Assay Kit (Cat. Q32855) and measured with the Invitrogen™ Qubit Flex™ Fluorometer. The TruSeq Stranded mRNA libraries (Cat. 20020594, Illumina, San Diego, CA, USA) were prepared using 100ng of total RNA per samples, using custom 384 xGen UDI-UMI adapters from IDT. The obtained libraries were then validated using the Fragment Analyzer system dsDNA 910 Reagent Kit (35–1500 bp) (Cat. DNF-910-K1000, Agilent) and with Qubit dsDNA HS Assay Kit (Cat. Q32854, Invitrogen). These were subsequently pooled equimolarly and sequenced on a Nextseq2000 (Illumina) by using a P2 flowcell with 50 base pairs paired-end reads, yielding on average 20 million reads per sample. Quality control of the sequence reads obtained from the FASTQ raw files was done using the FastQC (v0.11.8). Trimming of reads according to their quality and adapter presence was performed using TrimGalore (v0.6.5), and the resulting quality was once again assessed with FastQC (v0.11.8). SortMeRNA (v4.3.3) was used to filter out the rRNA reads and the resulting reads were subsequently aligned with the STAR (v2.7.3a) aligner to the reference genome fasta (GCA_000001405.15_GRCh38_no_alt_analysis_set.fna). Additional follow ups to the mapped (bam) files were carried out using PreSeq (v2.0.3), Sambamba (v0.7.0) and RSeQC (v3.0.1), after which read counts were generated with the Subread FeatureCounts module (v2.0.0) using the Homo_sapiens.GRCh38.106.ncbi.gtf file as an annotation. Normalization was achieved using the R-package edgeR (v3.28). Mean read counts obtained from this RNA sequencing can be found in an online database available at: https://esbl.nhlbi.nih.gov/Databases/KSBP2/.

Data analysis was performed using the integrated Differential Expression and Pathway analysis (iDEP) [16] and Kyoto Encyclopedia of Genes and Genomes (KEGG) [17] were used for enrichment analysis. Differentially expressed genes were analyzed with DESeq2. Upon comparing each condition to the untreated group, a minimum expression threshold of 2 reads per library across the replicates was used.

### 2.9 ciPTEC-OAT1 exposure to GPCRs inhibitors

To determine whether SCFAs exerted their effect via the activation of the GPCRs: GPR41, GPR43 and GPR109A, ciPTEC-OAT1 were co-incubated with SCFAs and the corresponding GPCR antagonist. Briefly, CiPTEC-OAT1 were cultured in 24 well-plates and upon complete maturation were exposed for 24 h to either acetate (1 mM), propionate (1 mM) or butyrate (1 mM), individually co-incubated with sodium β-hydroxybutyrate (SHB) (5 mM), a selective inhibitor for GPR41, GLPG0794 (0.1 µM) selective for GPR43 and mepenzolate bromide (100 µM) selective for GPR109A, respectively. To assess for OAT1 activity, the aforementioned fluorescein assay was carried out.

### 2.10 Histone deacetylase activity assay

HDAC enzyme activity was detected with a fluorometric HDAC commercially available kit (ab156064, Abcam, Cambridge, UK) according to manufacturer’s protocol. Briefly, cellular lysates were collected after 24 h exposure to each condition. TSA inhibitor, a class I and Class II HDAC inhibitors, was used as negative control. Fluorescence was monitored every 2 min over 30 min at 37°C using a GloMax™ Discover Microplate Reader (Promega, Leiden, The Netherlands) at emission wavelength of 415-445 nm and excitation wavelength of 365 nm. AFU data were normalized to their respective protein amount which was quantified using the Pierce™ Coomassie Plus (Bradford) Assay Kit (Thermo Fisher). Briefly, cell lysate and BCA standards (diluted in the HDAC assay extraction buffer) were loaded in a 96 well plate and incubated for 10 min at room temperature together with the Coomassie Plus Reagent. Protein amount was quantified using the GloMax™ Discover Microplate Reader (Promega, Leiden, The Netherlands) at absorbance 595nm. Background values were subtracted from the arbitrary absorbance unit and were normalized to the samples protein amount.

### 2.11 Statistical analysis

All statistical analyses were performed using GraphPad Prism (version 9.3.0, La Jolla, CA, USA). Prior to statistical analysis, the normality distribution of the data was tested using the Shapiro-Wilk test and based on data distribution one-way ANOVA or the non-parametric Kruskal-Wallis test were used. Multiple comparison adjustment was applied upon comparison among the untreated group, unless stated otherwise, to all tested conditions with either Dunnett’s or Dunn’s test as appropriate. All data are expressed as mean ± SEM and a *p* value < 0.05 was considered statistically significant.

## 3. RESULTS

### 3.1 SCFAs exposure boosts OAT1 activity via enhanced *SLC22A6* transcription

We first evaluated the safety of acetate, propionate and butyrate on proximal epithelial tubule cells by exposing ciPTEC-OAT1 to concentrations of 1-5 mM of the SCFAs for 24 h (Supplementary Fig. 1). Acetate appeared to be best tolerated, while butyrate was more toxic. Exposure to the SCFAs mix resulted in a comparable response as for butyrate alone. In light of these findings, we continued the current study with a concentration of 1 mM for each of the SCFAs, whether used alone or in combination.

Next, gene expression analysis *via* qRT PCR revealed that SCFAs increased the mRNA expression of *SLC22A6* (Fig. 1A). Notably, propionate and butyrate exposure yielded the highest increase in gene expression, whereas no synergistic effect was observed for the mix which was similar to the effect mediated by butyrate or propionate alone. In contrast, exposure to acetate alone did not increase *SLC22A6* gene expression compared to the untreated control. The increase in gene expression observed with butyrate and propionate was accompanied by an increase in protein production (Fig. 1B) and, more importantly, in OAT1 function (Fig. 1C). Among the SCFAs, butyrate exerted the most pronounced effect with a 71% increase in transport activity compared to the untreated control, a result that was also observed for the SCFAs mix. To validate the specificity of the observed enhancement in OAT1 activity, probenecid, an OAT1 substrate inhibitor, served as a negative control. Probenecid resulted in a substantial decrease in OAT1 activity down to 20%. Additional data on the effect of probenecid co-incubated with each SCFAs can be found in the supplementary information (Supplementary figure 2). Our results demonstrate that butyrate and propionate increased OAT1 activity.

**Figure 1.**
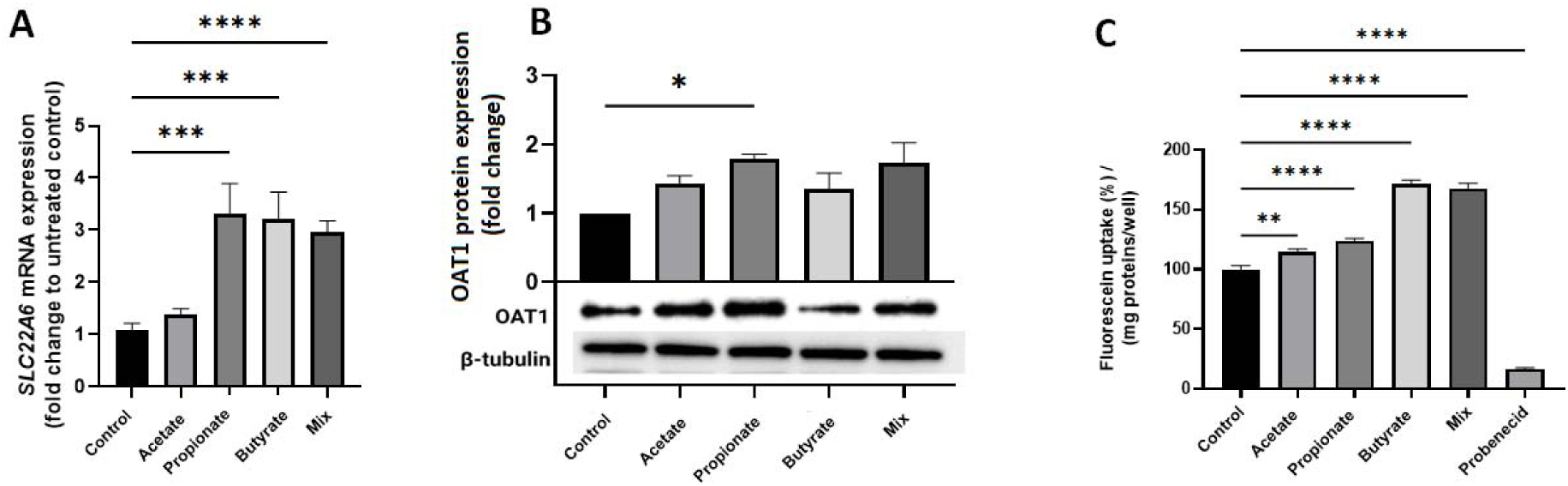
Exposure to SCFAs enhances *SLC22A6* transcription and OAT1 translation resulting in an enhanced OAT1 functionality. **A)** Gene expression of *SLC22A6* encoding for OAT1. **B)** OAT1 protein production and the respective representative Western blots. Protein expression levels were normalized against the housekeeping protein β-tubulin. **C)** Activity of the OAT1 transporter. All data are representative of 3-4 independent experiments and are reported as mean ± SEM. Data were checked for normal distribution with Shapiro-Wilk test and statistical significance was analyzed with One-Way ANOVA or Kruskal-Wallis test as appropriate. (\**p* < 0.05, \*\**p* < 0.01, \*\*\**p* < 0.001, \*\*\*\**p* < 0,0001).

### 3.2 Butyrate enhances the removal of IS in a kidney-on-chip system

To further assess the ability of SCFAs to enhance the removal of microbiota-generated uremic toxins, a KoC system, consisting of a bioengineered kidney tubule featuring an integrated basolateral flow representative of the systemic circulation and an apical compartment representing the tubular lumen compartment, was used (Fig. 2A) [18, 19]. Acetate, propionate and butyrate were added either alone or in combination to the basolateral compartment. After 24h exposure, the transepithelial transport of IS from the basolateral to the apical side was studied, representative of the movement of uremic toxins from blood to urine. Bioengineered kidney tubules treated with SCFAs showed an increased trend in IS excretion compared to the untreated control (Fig. 2B), with butyrate yielding the most pronounced increase in IS excretion. Surprisingly the SCFAs mix led to a somewhat decreased active transport (Fig. 2C). FITC-inulin assay was performed alongside to assess the monolayer integrity of the bioengineered kidney tubules, which appeared to be stable for all individual SCFAs tested when compared to the no treatment control, confirming that the apical IS concentrations reflected transport across the epithelial barrier.

**Figure 2:**
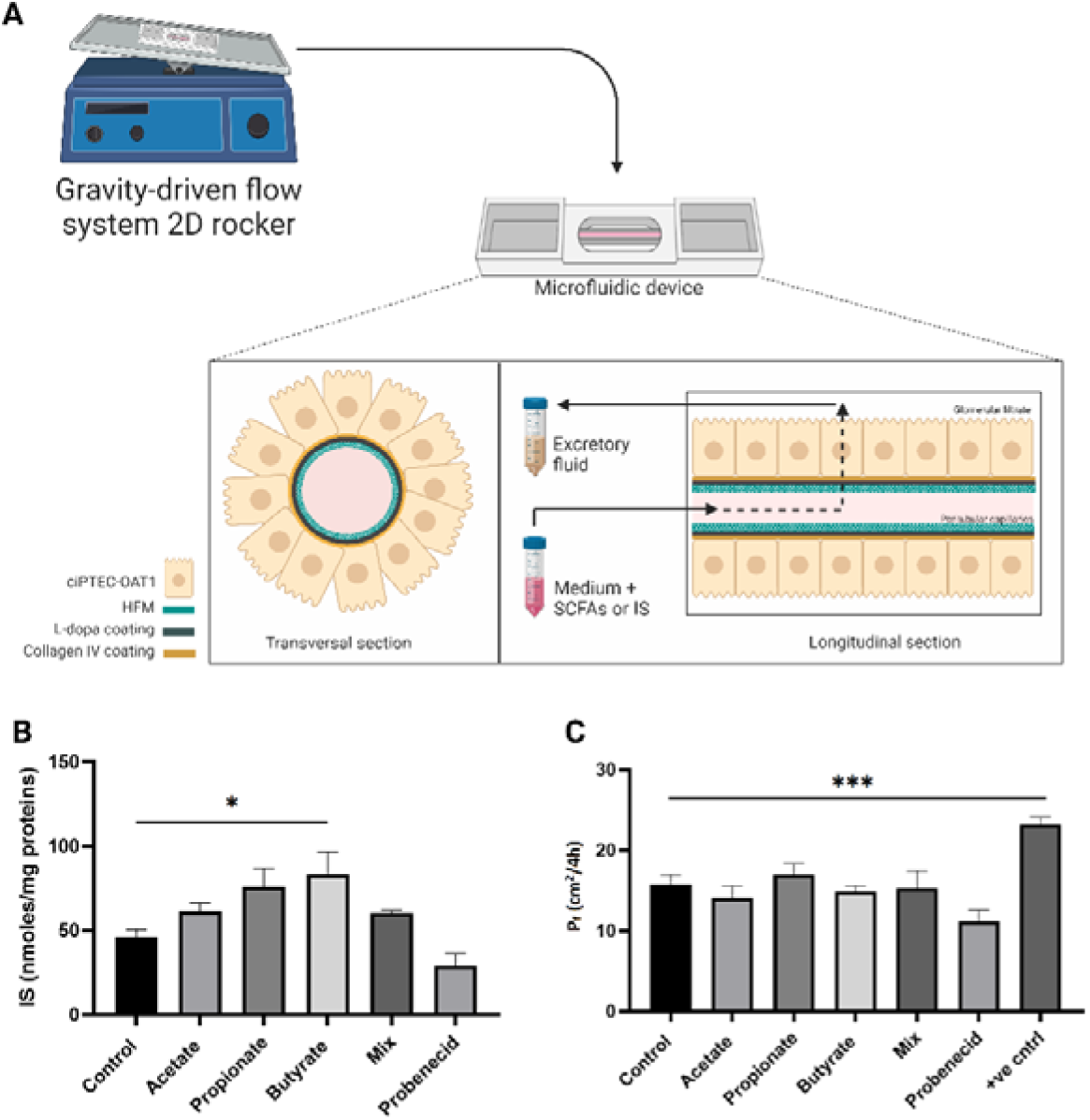
SCFAs enhance the removal of IS on a KoC system. **A)** Schematic representation of the experimental approach using the bioengineered kidney tubules in a perfusion system, created with Biorender. **B)** Removal of IS from the basolateral compartment towards the apical compartment of the KoC system. Data is normalized to the protein amount of each kidney tubule and reported as nmoles/mg proteins. **C)** FITC-inulin leakage as a measure of monolayer integrity of ciPTEC-OAT1 on the HFM. Negative control consisted of unseeded HFM reflecting for full leakage. Data are representative of 5 independent experiments. All data is reported as mean ± SEM. Data were analyzed with One-Way ANOVA and Dunnet’s test for multiple comparison adjustment; \**p* < 0.05, \*\*\**p* < 0.001.

### 3.3 The SCFA’s mediated increase in OAT1 activity is independent of GPCR signaling

Gene expression analysis was performed to assess whether exposure to SCFAs influenced mRNA expression of *FFAR3* encoding GPR41, *FFAR2* encoding GPR43 and *HCAR2* encoding GPR109A, known to be involved in SCFAs sensing. No significant changes were observed for any of the tested GPCRs genes, with mRNA levels being quite stable and similar to the untreated group (Fig. 3A). In agreement, no change in protein production was observed as levels of GPR41, GPR43 and GPR109A were below the level of detection (data not shown). Despite this, we investigated whether the observed increase in OAT1 activity was indirectly linked to GPCRs activation. Overall, we noticed that none of the inhibitors were able to block the functional boost of OAT1 observed following exposure to each of the SCFAs, further confirming that signaling of SFCAs to OAT1 observed here is independent of the GPCRs (Fig. 3B-D).

**Figure 3.**
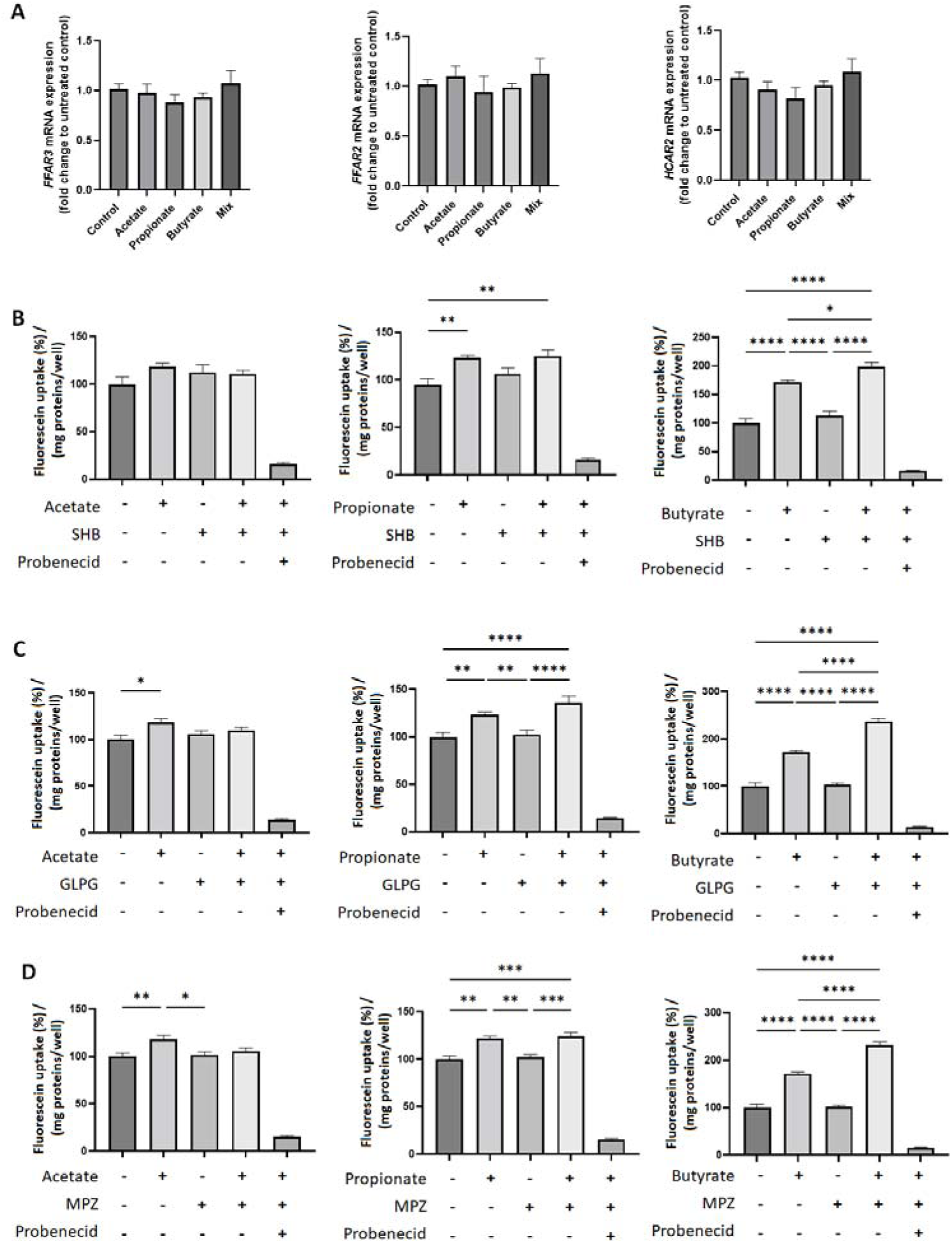
GPCRs do not show involvement in the SCFA’s mediated increase in OAT1 activity. **A)** Gene expression of the genes *FFAR3*, *FFAR2*, *HCAR2* encoding for GPR41, GPR43 and GPR109A respectively upon exposure to 1 mM acetate, propionate and butyrate either alone or in combination for 24 h. **B-D)** Activity of the OAT1 transporter upon a 24 h co-exposure with each SCFAs and the GPCRs inhibitors, namely GLPG074 a selective inhibitor for GPR41, SHB selective for GPR43 and mepenzolate selective for GPR109A. Data are representative of 3-4 independent experiments. All data is reported as mean ± SEM. Data were checked for normal distribution with Shapiro-Wilk test and statistical significance was analyzed with One-Way ANOVA or non-parametric Kruskal-Wallis test and adjusted for multiple comparison with either Dunnet’s or Dunn’s test as appropriate; \**p* < 0.05, ** *p* < 0.01, *** *p* < 0.001, **** *p* < 0.0001.

### 3.4 Transcriptomics analysis uncovers the ability of SCFAs to induce prominent and unique patterns of differentially expressed genes in proximal tubule cells

To further assess the overall effect of the SCFAs on our model, the gene expression signatures were evaluated. A total of 18376 genes passed the specified expression threshold of at least 0.5 counts per million (CPM) in at least one sample per each tested treatment. The Principal Component Analysis (PCA) plot shown in Fig. 4A displays each individual sample in a 2D space. The 86% variance observed in principal component 1 (PC1), indicates that the main source of variation in gene expression is linked to the treatment group; whereas the total variance recorded for the samples belonging to the same treatment group only accounted for 4% in variability, as depicted in PC2. Moreover, the PCA reports a close proximity between the untreated control and acetate as well as between butyrate and the mix. Propionate, on the other hand, does not cluster with neither of the other treatments nor the untreated control. These observed patterns of clusters are further reflected within the generated heatmap displaying the 1000 most variable genes observed in our samples (fig. 4B). Overall, all treatments induced both prominent and unique patterns of genes up- and down-regulations. Upon comparison of the untreated control to each of the treatments, a total of 19527 passed the expression filter of CPM > 0.5 in at least 2 samples. For this analysis we used a fold change (FC) > 2 and a false discovery rate (FDR) cutoff < 0.05 to define differentially expressed genes (DEG). Accordingly, among the identified expressed genes, acetate induced the upregulation and downregulation of only 1 gene when compared to the untreated control (fig. 4C), highlighting an almost identical transcript of gene expression patterns between both groups. When comparing propionate to the untreated control, a total of 987 genes were differentially expressed (822 upregulated, 165 downregulated). Butyrate and the mix, upon comparison to the untreated control yielded similar results, with a total of 3630 differentially expressed genes (2401 upregulated, 1229 downregulated) for butyrate, and 4034 differentially expressed genes (2613 upregulated, 1421 downregulated) for the mix, providing evidence that the mix of SCFAs had the biggest effect on transcription of gene expression. The Venn diagram in supplementary figure 3 displays in details the number of up- and down-regulated genes that are uniquely or shared between each SCFAs.

**Figure 4:**
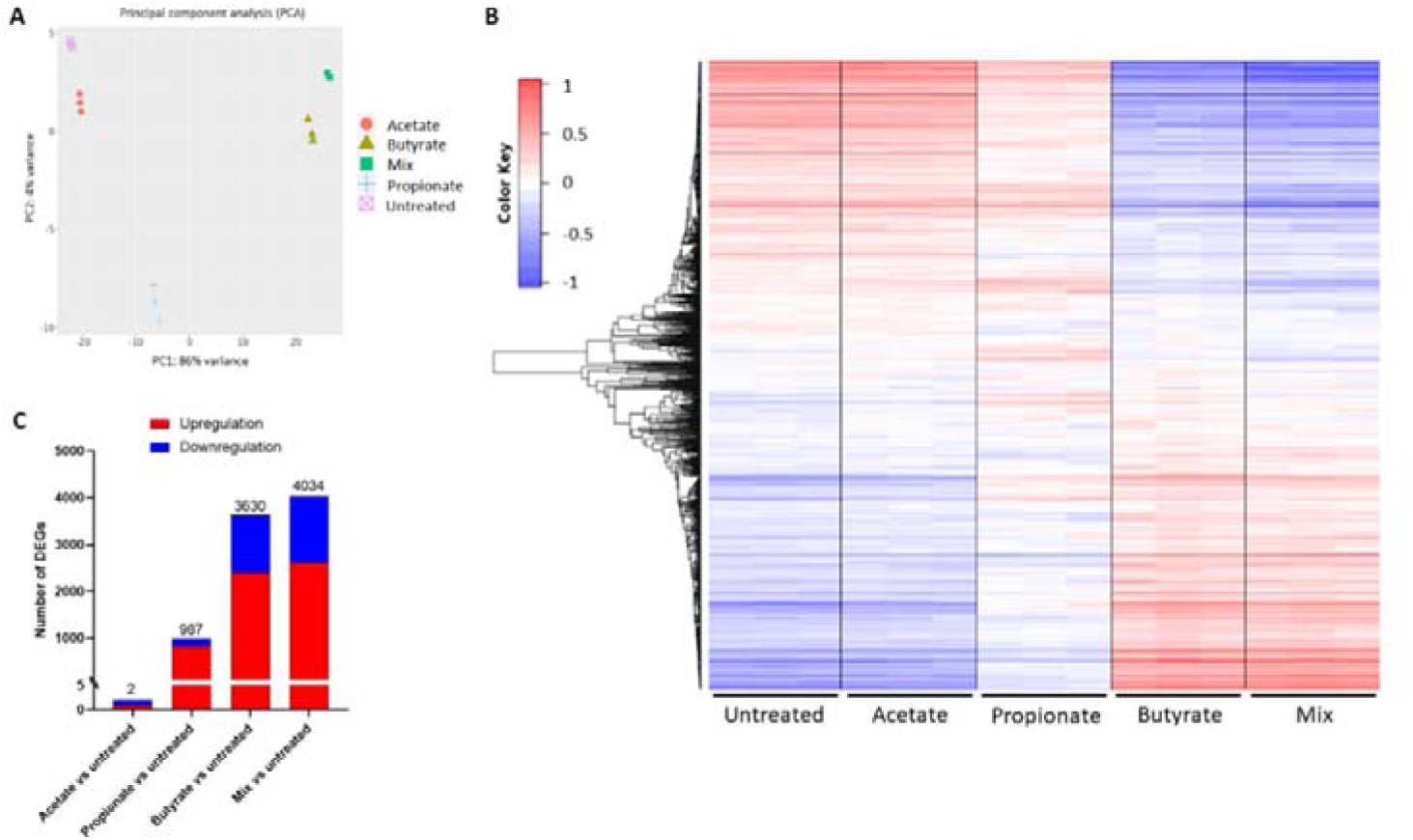
Gene expression analysis by RNA sequencing in ciPTEC-OAT1 exposed to SCFAs. **A)** PCA plot displaying the variance of 3 biological replicates per untreated control, acetate, propionate, butyrate and SCFAs mix samples. Each condition is represented by a distinct shape and color, corresponding to a 24 h exposure to 1 mM of acetate, propionate, butyrate either alone or in combination. **B)** Heatmap of the top 1000 differentially expressed genes among the different treatment groups. **C)** Number of statistically significant (FC > 2, FDR < 0.05) upregulated and downregulated DEGs upon comparison of each treatment group with the untreated control: acetate vs untreated group; propionate vs untreated group; butyrate vs untreated group; mix vs untreated group.

### 3.5 SCFAs regulate OATs through the cAMP signaling pathway

Transcriptomic analysis revealed that the activation of the cyclic AMP (cAMP) signaling pathway plays a pivotal step in the observed SCFAs mediated effects (figure 5A). In particular, propionate, butyrate and the SCFAs mix demonstrated to influence expression of genes encoding for adenylate cyclases (AD), mainly adenylate cyclase type 3 (*ADCY3*) and 9 (*ADCY9*) (figure 5B). The SCFAs mix also significantly upregulated the genes encoding for protein kinase A (PKA), namely the protein kinase cAMP-activated catalytic subunit alpha (*PRKACA*) (figure 5C), followed by an upregulation trend mediated also by butyrate and propionate. Upregulation of PKA, subsequently, leads to the modulation of a series of downstream signaling pathways as a result of their mediated phosphorylation activity, which upregulate the cAMP response element-binding protein 1 (*CREB1*) (figure 4D), culminating in the upregulation of target genes.

**Figure 5:**
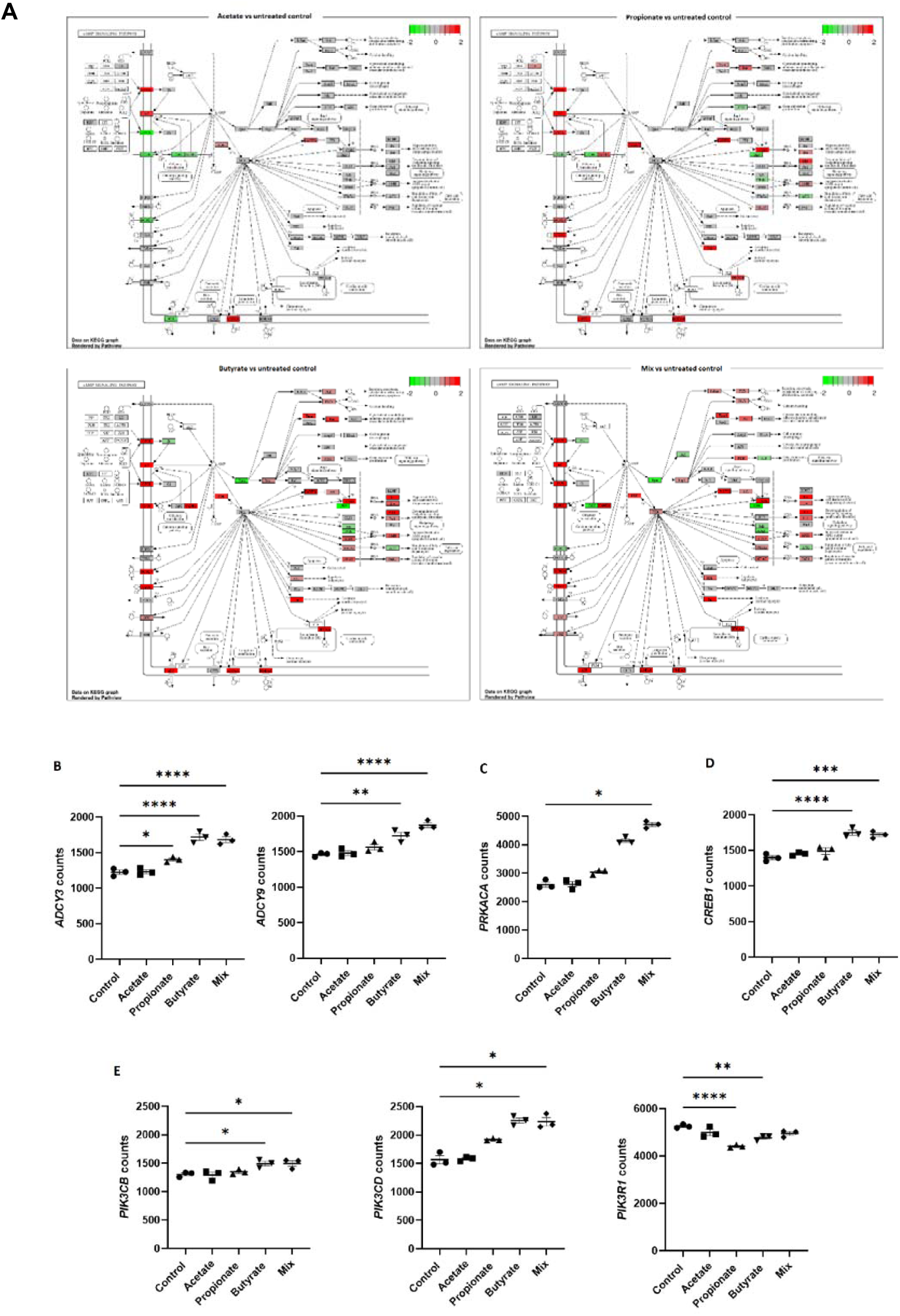
SCFAs regulate cAMP signaling pathway markers. **A)** Schematic representation of the cAMP signaling pathway upon exposure to acetate, propionate, butyrate or mix compared to the untreated control. Green: gene downregulation; red: gene upregulation. **B-E)** Gene expression of relevant markers within the cAMP signaling pathway as determined through RNA sequencing: **B**) adenyate cyclases (*ADCY3* and *ADCY9*); **C**) protein kinases (*PRKACA*); **D**) cAMP response element-binding protein 1 (*CREB1*); **E**) phosphoinositide 3 kinase class 1A (*PIK3CB, PIK3CD* and *PIK3R1*). Data are reported as mean ± SEM. Data were checked for normal distribution with Shapiro-Wilk test and statistical significance was analyzed with One-Way ANOVA or Kruskal-Wallis test as appropriate. (\**p* < 0.05, \*\**p* < 0.01, \*\*\**p* < 0.001, \*\*\*\**p* < 0.0001).

Furthermore, upregulation of *ADCY3* and *ADCY9* mediated by butyrate and the SCFAs mix leads to a series of downstream pathways able to upregulate gene expression of the intracellular signal transducer phosphoinositide 3 kinase (*PI3K*) class 1A, mainly PI3K catalytic subunit B (*PIK3CB*) and D (*PIK3CD*) (figure 5E), initiating a signaling cascade within the PI3K-Akt signaling pathways (supplementary figure 4), which has previously been implicated in OAT1 signaling [20]. Propionate, on the other hand, inhibited PI3K regulatory subunit 1 (*PIK3R1*), an effect that was also mediated by butyrate and followed with a trend by the SCFAs mix.

### 3.6 SCFAs influence the expression of genes encoding for histone deacetylase and inhibit their activity in ciPTEC-OAT1

Given that SCFAs are well-established inducers of HDAC inhibition, we analyzed the effects of individual SCFAs and their combination on the expression of genes encoding for HDAC enzymes. The expression levels of all 18 identified HDAC genes are provided as log2-fold change in the supplementary figure 5 heatmap, providing insight into the extent of modulation induced by each SCFAs. Statistically significant changes in gene expression are shown in figure 6A-B as gene counts. Exposure to butyrate and the SCFAs mix elicited the most potent effects, followed by a comparatively milder effect induced by propionate, while acetate did not yield significant changes in gene expression. Overall, the observed differential expression patterns for each SCFA remained consistent, with no treatment yielding opposing effects in gene regulation compared to the others. Among the 12 genes that were significantly differentially expressed, those belonging to class I (*HDAC1*, *HDAC2* and *HDAC3*) and most genes in class III (*SIRT2*, *SIRT4* and *SIRT7*) exhibited upregulation. On the other hand, most genes belonging to class II (*HDAC6*, *HDAC7* and *HDAC9*) yielded downregulation. Notably, *HDAC7* yielded the most pronounced downregulation, with gene expression inhibition exceeding 50% in gene counts following exposure to butyrate and the SCFAs mix. Downregulation was also observed for some class III genes, including *SIRT3* and *SIRT7*. Overall, the highest gene counts levels were reported for *HDAC1* and *HDAC7*.

**Figure 6:**
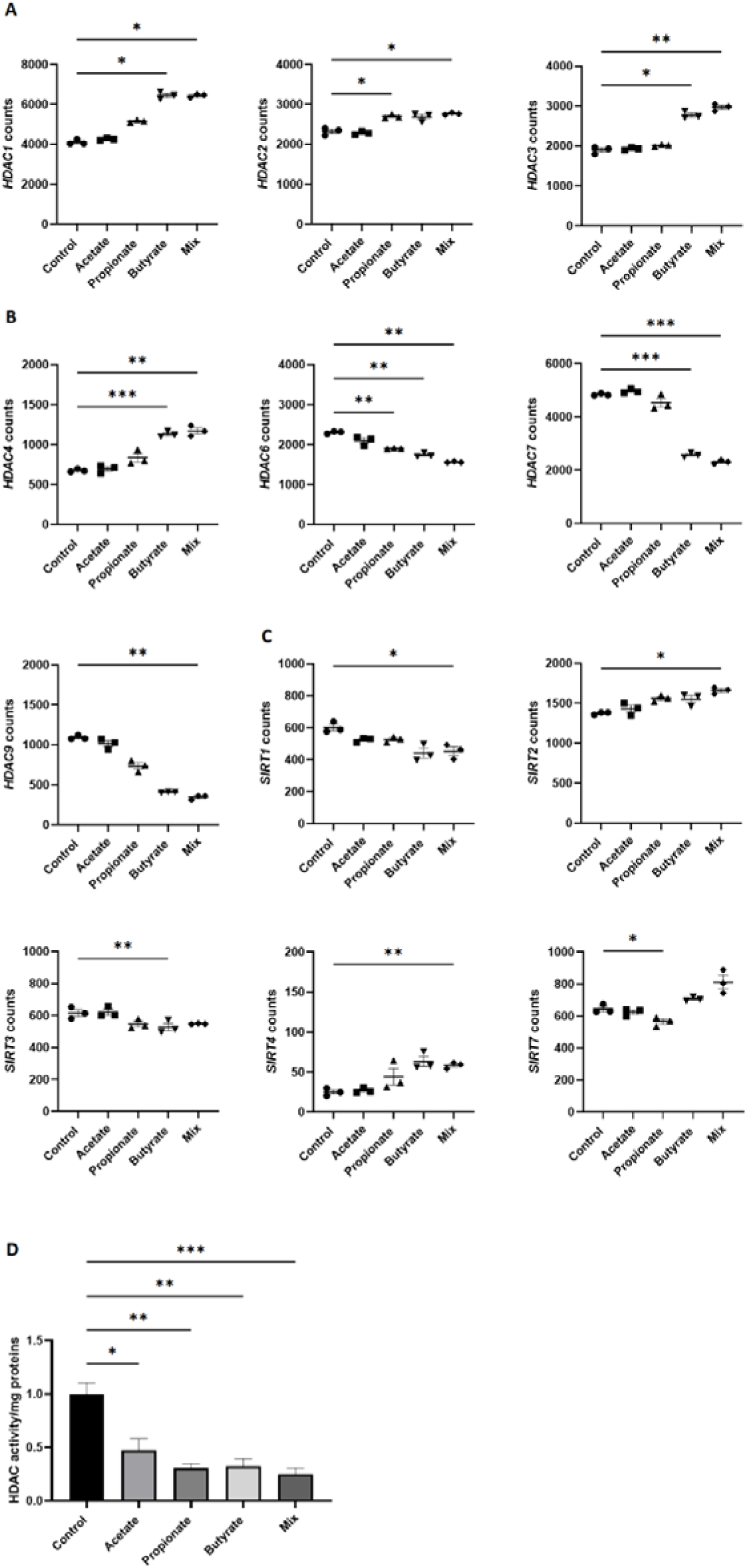
SCFAs influence the expression of HDAC genes. Counts of: **A)** Class I HDAC genes: *HDAC1*, *HDAC2* and *HDAC3*; **B)** Class II: *HDAC4*, *HDAC6*, *HDAC7* and *HDAC9*; **C)** Class III: *SIRT1, SIRT2, SIRT3, SIRT4* and *SIRT7*. Data is reported as mean ± SEM. Data were checked for normal distribution with Shapiro-Wilk test and statistical significance was analyzed with One-Way ANOVA or non-parametric Friedman test and adjusted for multiple comparison with either Dunnet’s or Dunn’s test as appropriate; \**p* < 0,05, \*\**p* < 0,01, \*\*\**p* < 0,001, \*\*\*\**p* < 0,0001. D) SCFAs effect on HDAC activity inhibition following 24 h exposure to acetate, propionate, and butyrate either alone or in combination. Statistical significance was analyzed with Kruskal-Wallis test and Dunn’s test for multiple comparison adjustment; \**p* < 0,05, \*\**p* < 0,01, *** *p* < 0,001, \*\*\*\**p* < 0,0001.

Furthermore, we investigated whether the observed increase in OAT1 expression and activity could instead be dependent upon the ability of SCFAs to inhibit HDAC activity (figure 6C). Our results demonstrate that all SCFAs can effectively inhibit HDAC activity in proximal tubule cells. Particularly, the SCFAs mix had the strongest inhibition when compared to the untreated control.

### 3.7 Uptake and efflux transporters gene expression is affected upon SCFAs exposure

Finally, we evaluated the impact of SCFAs on the expression levels of transporters involved in renal tubular waste excretion in addition to *SLC22A6* (figure 7). Consistent with our qRT PCR findings, transcription of *SLC22A6* was significantly upregulated by propionate, butyrate and the SCFAs mix treatments, however, this was not observed for *SLCO4C1* (figure 7B). Influx transporters for metabolic waste, including IS, work in concerted action with efflux transporters, of which a strong upregulation in *ABCG2* and *ABCB1* was observed (figure 7C) [19, 21]. Specifically, butyrate upregulated *ABCG2*, encoding BCRP, while the SCFAs mix treatment predominantly upregulated *ABCB1*, encoding P-glycoprotein. On the other hand, transcription of *SLC47A2*, encoding MATE2K, was strongly downregulated by both butyrate and the mix treatment.

**Figure 7:**
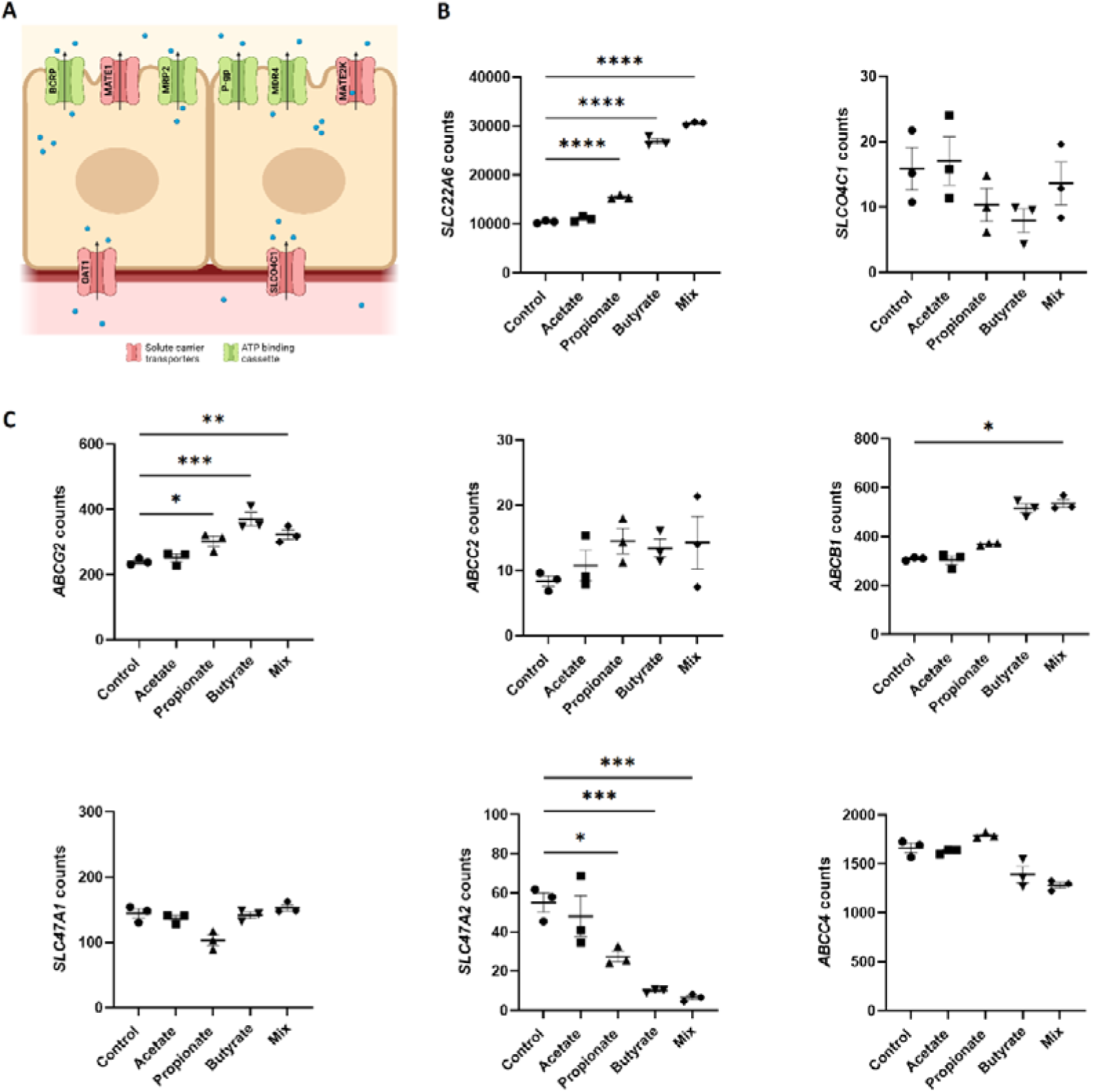
Uptake and efflux transporters gene expression is affected upon SCFAs exposure. A) schematic representation of apical and basal transporters. B) Counts of ciPTEC-OAT1 genes encoding for drug transporters: influx transporters: OAT1/*SLC22A6*; OATP4C1/*SLCO4C1*. C) efflux transporters: BCRP/*ABCG2*; MRP2/*ABCC2*; MDR1/*ABCB1*; MATE1/*SLC47A1*; MATE2K/*SLC47A2*; MRP4/*ABCC4*. CPM > 0.5, FC > 2 and FDR cutoff < 0.05. All data is reported as mean ± SEM. Data were checked for normal distribution with Shapiro-Wilk test and statistical significance was analyzed with One-Way ANOVA or non-parametric Friedman test and adjusted for multiple comparison with either Dunnet’s or Dunn’s test as appropriate; \**p* < 0,05, \*\**p* < 0,01, \*\*\**p* < 0,001, \*\*\*\**p* < 0,0001.

## 4. DISCUSSION

The data presented in this study demonstrates that SCFAs enhance the activity of OAT1 in kidney proximal tubule cells by inhibiting HDAC enzyme activity and modulating the cAMP signaling pathway. Our findings confirm that ciPTEC-OAT1 can safely tolerate SCFAs concentrations up to 1 mM for 24 h, with this exposure significantly inducing OAT1 expression and activity compared to untreated cells. Interestingly, SCFAs did not mediate their effects via the activation or upregulation of GPR41, GPR43 and GPR109A.

OAT1 plays a critical role in the secretion of metabolic waste excretion, including uremic toxins, thereby preserving kidney homeostasis. Upon development and progression of kidney disease, OAT1 function becomes impaired, contributing to the development of uremic syndrome and associated complications [22, 23]. Several studies have established a correlation between OAT1 downregulation and the deterioration of renal function following kidney injury both clinically and *in vivo*, highlighting the role played by OAT1 in interorgan communication and implications in kidney disease [22, 24, 25]. Here, we provide evidence that SCFAs can directly modulate OAT1 function, highlighting their potential therapeutic value.

SCFAs are known to exert nephroprotective effects in vivo, improving kidney function after AKI by mitigating the inflammatory response, reducing oxidative stress and acting as histone deacetylase inhibitors [26–31]. Although acetate is the most abundant SCFA in the systemic circulation [32], our observations revealed that propionate and butyrate exert the most potent effects on OAT1 activity in our kidney cell line. Specifically, butyrate exhibited the strongest regulation on OAT1 activity, while propionate on OAT1 gene transcription and translation. Both SCFAs were observed to improve renal biochemical indices/parameters in murine models of renal failure, with butyrate being proposed to mitigate kidney fibrosis by decreasing the serum concentrations of the uremic toxin TMAO [29, 30]. Accordingly, upon evaluating the physiological relevance of our findings on the enhanced OAT1 activity, we observed an increase in the butyrate-mediated secretory clearance of IS during the perfusion of the bioengineered kidney tubules with IS after exposure to SCFAs. Nevertheless, the mechanisms by which SCFAs improve kidney function remain to be fully elucidated. The use of KoC systems offers an opportunity for prolonged *in vitro* exposure studies to further explore SCFAs

Upon evaluating the cAMP signaling pathway following SCFAs exposure, we observed an upregulation mediated by butyrate and the SCFAs mix of genes encoding for adenylate cyclases (*ADCY3* and *ADCY9)*. Consequently, this led to the activation of cAMP and PKA (*PRKACA*) genes upregulation, which in turn regulated *CREB1*transcriptional activity. *CREB1* serves as a key factor involved in both constitutive and inducible transcriptional regulation, with the latter being modulated by the extracellular activation of adenylate cyclase and PKA. Several studies have documented the ability of butyrate to activate the cAMP-PKA-CREB signaling pathway in various cell types, including intestinal [33], neuronal [34] and immune cells lines [35]. Particularly, Wang *et al.* elucidated a potential mechanism for this pathway activation, attributing it to the ability of butyrate to increase intracellular ATP levels; and that acetate and propionate might similarly activate the pathways due to their ATP-generating capabilities [33]. Accordingly, the extent of pathway activation by these SCFAs may thus vary across cell types according to the expression of acetyl-CoA, propionyl-CoA and butyryl-CoA synthetases, which regulate ATP production from acetate, propionate and butyrate, respectively [33]. The involvement of CREB1 in the constitutive transcriptional regulation of an OAT family member was demonstrated by the induction of the expression of *SLC22A8* encoding OAT3 via the PKA-mediated phosphorylation of CREB1 [36]. Wang *et al.* further investigated this PKA-mediated transcriptional regulation showing that PKA activation enhanced the expression and transport activity of OAT3, while prolonged PKA activation reduced the transporter’s degradation [37]. Although these studies solely focused on OAT3, the genomic clustering of the OATs genes argues for a coordinated regulation for *SLC22A6* transcription by the same signaling pathways [20, 38].

Further investigation into the downstream signaling pathways revealed that SCFAs modulate the PI3K-Akt pathway (supplementary Fig 4). PI3K enzymes play a pivotal role in regulating several intracellular processes, including gene expression, through the phosphorylation of phosphatidylinositol, a signal transduction precursor of several second messenger molecules [39]. Our findings identified both an up- and down-regulation of PI3K genes expression being mediated by butyrate and the SCFAs mix, reflecting the dual role of butyrate in modulating this pathway. While some studies have highlighted butyrate’s potential to enhance PI3K signaling via the PI3K pathway [40, 41], others have demonstrated its inhibitory effects [42, 43]. The indirect involvement of PI3K in OAT1 regulation has previously been reported by Caetano-Pinto et al., with EGF-induced PI3K activation influencing cAMP production and PKA activity, thereby stimulating OAT function [44][20, 37]. Previous studies suggested that both OAT1 and OAT3 expression may be regulated by both the PI3K and cAMP downstream signaling pathways, with a possible influence from SUMOylation activity due to SCFAs [45]. Moreover, our findings align with the theory of the kidneys ability to “remote sense and signal” elevated levels of metabolites, such as IS, through pathways like epidermal growth factor receptor (EGFR) activation, ultimately leading to the activation of OAT1-mediated secretory function [8, 46] [47]. It is relevant to note that the expression levels of *SLC22A6* were approximately 10’000 times higher than those of all other transporters. This disparity in expression levels is attributable to our use of the transfected cell line overexpressing the transporter itself which equals the expression levels in native human kidneys [48].

SCFAs have been recognized to exert their effects through the activation of GPR41, GPR43 and GP109A [30, 49–53], nevertheless our results indicate that their effects on OAT1 function are independent of these receptors. Co-incubation with selective inhibitors for these GPCRs did not diminish the enhanced OAT1 activity, nor did SCFAs significantly alter the gene expression of these receptors. This is consistent with previous studies reporting that SCFA-mediated effects can occur independently of GPCR activation [54–56] and is likely due to differences in GPCR expression between tissues, with kidney expression being relatively low [57].

Most likely, the SCFAs mediate their effect in the kidneys through direct nuclear HDAC inhibition. This is in agreement with findings reported for the activation of the Aryl hydrocarbon receptor (AhR), a uremic toxins receptor [58], for which the chromatin conformation in the promoter region that facilitate the binding of the AhR-ligand complex has been found to be correlated with the ability of butyrate to act as an HDAC inhibitor [59]. Our results revealed a significant inhibitory activity of each tested SCFA on HDACs, in line with previous studies that highlighted the role of HDAC inhibition in attenuating kidney fibrosis [60–62] and thus improving kidney function in CKD. Accordingly, to further explore HDAC inhibition, we assessed the expression levels of all 18 identified HDAC genes. Overall, we observed that genes encoding for class II HDACs, exhibited the most significant downregulation compared to other classes, suggesting that these HDACs might represent the main target of SCFAs. Class II HDACs can translocate between the cytosol and nucleus [63]. Expression of HDAC6 and HDAC9 have been implicated in kidney disease pathogenesis, with HDAC6 contributing to injury observed in cisplatin-induced AKI [64–66] and HDAC9 associated with kidney fibrosis [67], podocytes injury [68] and vascular calcification [69, 70]. While HDAC7 yielded the strongest SCFAs-mediated inhibition in our results, the role of HDAC7 remains less defined, hence it has been suggested to be involved in sodium reabsorption [71]. Interestingly, Wang et al. found that histone acetylation was involved in the regulation of OAT2 in hepatocellular carcinoma cells. OAT2 was found to be downregulated due to hypoacetylation of H4K16ac in hepatocellular carcinoma cells, mediated by KAT8 and HDAC7 [72]. Although the exact mechanism by which these HDACs protect the kidneys from injury is not fully elucidated, it is hypothesized that it may partly involve their effect on *SLC22A6* gene expression. HDACs are known regulators of cellular responses to cAMP signaling, playing a role in the deacetylation of histones associated with cAMP response elements (CREs) to which CREB binds following phosphorylation [1, 73]. As a result, HDACs inhibition could enhance gene transcription of cAMP pathway-regulated genes, as we observed for *SLC22A6* [1, 73]. HDACs have also been found to be involved in the PI3K pathway, with HDAC6 playing a role in EGFR degradation [74]. Accordingly, the observed upregulation of EGFR (supplementary Fig. 6) may be the result of a reduced degradation facilitated by the downregulation of HDAC6 gene expression, which is thereby likely contributing to the upregulation of PI3K gene expression. Follow-up studies should focus on elucidating which specific HDAC gene(s) are involved in the observed effect on OAT1 expression, abundance and function in the kidney cell line.

HDAC inhibition affects the cAMP signaling pathway, modulating CREB1 and PI3K gene expressions. These regulate cell metabolism and stress responses, promotes proliferation and survival, enhance *SLC22A6*/OAT1 function, supporting overall cellular health, function, and resilience, thus enhancing cellular fitness. *CREB1* activation was shown to provide protection against oxidative stress induced by cellular damage and activate the expression of antioxidant genes [75]. Recent studies have highlighted the role of *CREB1* in yielding protection against kidney stones both *in vitro* and *in vivo*, possibly by alleviating the activation of inflammatory pathways, exerting anti-inflammatory, antioxidant and anti-apoptotic effects [76, 77]. Accordingly, CREB1 modulation could have therapeutic implications for podocytes injury, nephropathy and kidney ischemic reperfusion injury [78, 79]. The PI3K pathway was also shown to be involved in kidney cells metabolism regulation *in vivo*, improving kidney cells metabolism [80] and delaying kidney injury [81, 82]. PI3K is a known regulator of autophagy, and as such PI3K pathway modulation can confer protection to kidney function via inhibition of inflammatory reactions [83–85]. The involvement of PI3K pathway in kidney physiological regulation in response to stress, injury inflammation and apoptosis was also recently reported [86].

We acknowledge certain limitations within our study. Given that mRNA levels do not always accurately represent protein abundance and function, further experiments should evaluate the effects of SCFAs on HDAC enzyme as well as on the intricate signaling cascade by them initiated, which potentially involves numerous post-translational modifications. Overall, the interplay between SCFAs and OAT1 is multifaceted, and our understanding of their modulation of physiological processes remains limited.

To conclude, our study provides compelling evidence that propionate and butyrate significantly enhance OAT1 activity through *SLC22A6* gene upregulation. This effect is independent of GPCRs activation and rather involves class II HDAC enzymes inhibition, which modulates cAMP signaling pathways and downstream effectors, such as CREB1 and PI3K. These findings highlight the therapeutic potential of SCFAs as modulators of kidney function and underscore the need for further studies to explore their long-term benefits and applications in treating kidney diseases.

## CRediT authorship contribution statement

**Laura Giordano**: Conceptualization, Data curation, Formal analysis, Investigation, Methodology, Validation, Visualization, Writing – original draft, Writing – review & editing. **Sabbir Ahmed**: Investigation. **Thomas Van Der Made**: Supervision, Writing – review & editing. **Rosalinde Masereeuw**: Conceptualization, Funding acquisition, Supervision, Writing – review & editing. **Silvia Maria Mihaila**: Conceptualization, Supervision, Writing – review & editing.

## Acknowledgments

This work was supported by the EU Horizon 2020 research and innovation programme under the Marie SkłodowskaCurie grant agreement STRATEGY-CKD H2020-2019-ETN (860329), the Dutch Kidney Foundation Kolff+ program (22OK1019) and PPP Allowance made available by Health∼Holland, Top Sector Life Sciences & Health (LSHM20045-SGF) to the Association of Collaborating Health Foundations (SGF) to stimulate public-private partnerships. R.M. is a member of the ESAO/ERA-EDTA-endorsed Work Group EUTox.

## ABBREVIATIONS

OAT1: Organic anion transporter-1
IS: indoxyl sulfate
TMAO: trimethylamine N-oxide
SLC: solute carrier transporters
ABC: ATP-binding cassette
OAT3: Organic anion transporter-3
BCRP: breast cancer resistance protein
MRP2: multidrug resistance-associated proteins 2
MRP4: multidrug resistance-associated proteins 4
SCFAs: short chain fatty acids
KoC: Kidney-on-chip
ciPTEC-OAT1: conditionally immortalized proximal tubule epithelial cell line overexpressing OAT1
LDH: lactate-dehydrogenase
AFU: Arbitrary fluorescence unit
ATPs: ATP synthase
HFM: hollow-fiber membranes
CPM: counts per million
PCA: Principal Component Analysis
FC: fold change (
FDR: false discovery rate
DEG: differentially expressed genes
cAMP: cyclic AMP
AD: adenylate cyclases
*ADCY3*: adenylate cyclase type 3
*ADCY9*: adenylate cyclase type 9
PKA: protein kinase A
*PRKACA*: protein kinase cAMP-activated catalytic subunit alpha
*CREB1*: cAMP response element-binding protein 1
*PI3K*: phosphoinositide 3 kinase
*PIK3CB*: PI3K catalytic subunit B
*PIK3CD*: PI3K catalytic subunit D
EGF: Repidermal growth factor receptor
AhR: Aryl hydrocarbon receptor.

## References

1. Caetano-Pinto, P. and S.H. Stahl, Renal Organic Anion Transporters 1 and 3 In Vitro: Gone but Not Forgotten. Int J Mol Sci, 2023. 24(20).

2. Ermakov, V.S., J.C. Granados, and S.K. Nigam, Remote effects of kidney drug transporter OAT1 on gut microbiome composition and urate homeostasis. JCI Insight, 2023. 8(21).

3. Torday, J.S., Homeostasis as the Mechanism of Evolution. Biology, 2015. 4(3): p. 573–590.

4. Vaziri, N.D., et al., Chronic kidney disease alters intestinal microbial flora. Kidney Int, 2013. 83(2): p. 308–15.

5. Kanbay, M., et al., The crosstalk of gut microbiota and chronic kidney disease: role of inflammation, proteinuria, hypertension, and diabetes mellitus. International Urology and Nephrology, 2018. 50(8): p. 1453–1466.

6. Evenepoel, P., R. Poesen, and B. Meijers, The gut-kidney axis. Pediatr Nephrol, 2017. 32(11): p. 2005–2014.

7. Magliocca, G., et al., Short-Chain Fatty Acids in Chronic Kidney Disease: Focus on Inflammation and Oxidative Stress Regulation. Int J Mol Sci, 2022. 23(10).

8. Jansen, J., et al., Remote sensing and signaling in kidney proximal tubules stimulates gut microbiome-derived organic anion secretion. Proc Natl Acad Sci U S A, 2019. 116(32): p. 16105–16110.

9. van der Hee, B. and J.M. Wells, Microbial Regulation of Host Physiology by Short-chain Fatty Acids. Trends in Microbiology, 2021. 29(8): p. 700–712.

10. Nieskens, T.T., et al., A Human Renal Proximal Tubule Cell Line with Stable Organic Anion Transporter 1 and 3 Expression Predictive for Antiviral-Induced Toxicity. Aaps j, 2016. 18(2): p. 465–75.

11. Wilmer, M.J., et al., Novel conditionally immortalized human proximal tubule cell line expressing functional influx and efflux transporters. Cell Tissue Res, 2010. 339(2): p. 449–57.

12. Mihaila, S.M., et al., Drugs Commonly Applied to Kidney Patients May Compromise Renal Tubular Uremic Toxins Excretion. Toxins (Basel), 2020. 12(6).

13. Jansen, J., et al., Human proximal tubule epithelial cells cultured on hollow fibers: living membranes that actively transport organic cations. Sci Rep, 2015. 5: p. 16702.

14. Jochems, P.G.M., et al., A combined microphysiological-computational omics approach in dietary protein evaluation. NPJ Sci Food, 2020. 4(1): p. 22.

15. Ahmed, S., et al., A robust, accurate, sensitive LC-MS/MS method to measure indoxyl sulfate, validated for plasma and kidney cells. Biomed Chromatogr, 2022. 36(5): p. e5307.

16. Ge, S.X., E.W. Son, and R. Yao, iDEP: an integrated web application for differential expression and pathway analysis of RNA-Seq data. BMC Bioinformatics, 2018. 19(1): p. 534.

17. Kanehisa, M. and S. Goto, KEGG: kyoto encyclopedia of genes and genomes. Nucleic Acids Res, 2000. 28(1): p. 27–30.

18. Faria, J., et al., Bioengineered Kidney Tubules Efficiently Clear Uremic Toxins in Experimental Dialysis Conditions. Int J Mol Sci, 2023. 24(15).

19. Jansen, J., et al., Bioengineered kidney tubules efficiently excrete uremic toxins. Sci Rep, 2016. 6: p. 26715.

20. Pou Casellas, C., et al., Regulation of solute carriers oct2 and OAT1/3 in the kidney: a phylogenetic, ontogenetic, and cell dynamic perspective. Physiol Rev, 2022. 102(2): p. 993–1024.

21. Nigam, S.K., et al., The Systems Biology of Drug Metabolizing Enzymes and Transporters: Relevance to Quantitative Systems Pharmacology. Clin Pharmacol Ther, 2020. 108(1): p. 40–53.

22. Tan, S.P.F., et al., Effect of Chronic Kidney Disease on the Renal Secretion via Organic Anion Transporters 1/3: Implications for Physiologically-Based Pharmacokinetic Modeling and Dose Adjustment. Clin Pharmacol Ther, 2022. 112(3): p. 643–652.

23. Brandoni, A. and A.M. Torres, Altered Renal Expression of Relevant Clinical Drug Transporters in Different Models of Acute Uremia in Rats. Role of Urea Levels. Cellular Physiology and Biochemistry, 2015. 36(3): p. 907–916.

24. Bush, K.T., P. Singh, and S.K. Nigam, Gut-derived uremic toxin handling in vivo requires OAT-mediated tubular secretion in chronic kidney disease. JCI Insight, 2020. 5(7).

25. Sirijariyawat, K., et al., Impaired renal organic anion transport 1 (SLC22A6) and its regulation following acute myocardial infarction and reperfusion injury in rats. Biochimica et Biophysica Acta (BBA) - Molecular Basis of Disease, 2019. 1865(9): p. 2342–2355.

26. Andrade-Oliveira, V., et al., Gut Bacteria Products Prevent AKI Induced by Ischemia-Reperfusion. J Am Soc Nephrol, 2015. 26(8): p. 1877–88.

27. Machado, R.A., et al., Sodium butyrate decreases the activation of NF-κB reducing inflammation and oxidative damage in the kidney of rats subjected to contrast-induced nephropathy. Nephrol Dial Transplant, 2012. 27(8): p. 3136–40.

28. Sun, X., et al., Histone deacetylase inhibitor, sodium butyrate, attenuates gentamicin-induced nephrotoxicity by increasing prohibitin protein expression in rats. Eur J Pharmacol, 2013. 707(1-3): p. 147–54.

29. Wang, S., et al., Quantitative reduction in short-chain fatty acids, especially butyrate, contributes to the progression of chronic kidney disease. Clin Sci (Lond), 2019. 133(17): p. 1857–1870.

30. Mikami, D., et al., Short-chain fatty acid mitigates adenine-induced chronic kidney disease via FFA2 and FFA3 pathways. Biochimica et Biophysica Acta (BBA) - Molecular and Cell Biology of Lipids, 2020. 1865(6): p. 158666.

31. Marques, F.Z., et al., High-Fiber Diet and Acetate Supplementation Change the Gut Microbiota and Prevent the Development of Hypertension and Heart Failure in Hypertensive Mice. Circulation, 2017. 135(10): p. 964–977.

32. Lange, O., M. Proczko-Stepaniak, and A. Mika, Short-Chain Fatty Acids-A Product of the Microbiome and Its Participation in Two-Way Communication on the Microbiome-Host Mammal Line. Curr Obes Rep, 2023. 12(2): p. 108–126.

33. Wang, A., et al., Butyrate activates the cAMP-protein kinase A-cAMP response element-binding protein signaling pathway in Caco-2 cells. J Nutr, 2012. 142(1): p. 1–6.

34. DeCastro, M., et al., Short chain fatty acids regulate tyrosine hydroxylase gene expression through a cAMP-dependent signaling pathway. Molecular Brain Research, 2005. 142(1): p. 28–38.

35. Diakos, C., et al., Novel mode of interference with nuclear factor of activated T-cells regulation in T-cells by the bacterial metabolite n-butyrate. J Biol Chem, 2002. 277(27): p. 24243–51.

36. Ken, O., et al., Human Organic Anion Transporter 3 Gene Is Regulated Constitutively and Inducibly via a cAMP-Response Element. Journal of Pharmacology and Experimental Therapeutics, 2006. 319(1): p. 317.

37. Wang, H., J. Zhang, and G. You, Activation of Protein Kinase A Stimulates SUMOylation, Expression, and Transport Activity of Organic Anion Transporter 3. Aaps j, 2019. 21(2): p. 30.

38. Bhatnagar, V., et al., Analyses of 5’ regulatory region polymorphisms in human SLC22A6 (OAT1) and SLC22A8 (OAT3). J Hum Genet, 2006. 51(6): p. 575–580.

39. Golden, L.H. and K.L. Insogna, The expanding role of PI3-kinase in bone. Bone, 2004. 34(1): p. 3–12.

40. Tang, G., et al., Butyrate ameliorates skeletal muscle atrophy in diabetic nephropathy by enhancing gut barrier function and FFA2-mediated PI3K/Akt/mTOR signals. Br J Pharmacol, 2022. 179(1): p. 159–178.

41. Zhou, Z., et al., Sodium butyrate attenuated neuronal apoptosis via GPR41/Gβγ/PI3K/Akt pathway after MCAO in rats. Journal of Cerebral Blood Flow & Metabolism, 2020. 41: p. 267–281.

42. Mathew, O.P., et al., Cellular Effects of Butyrate on Vascular Smooth Muscle Cells are Mediated through Disparate Actions on Dual Targets, Histone Deacetylase (HDAC) Activity and PI3K/Akt Signaling Network. Int J Mol Sci, 2019. 20(12).

43. Sun, X., et al., Lactiplantibacillus plantarum NKK20 Increases Intestinal Butyrate Production and Inhibits Type 2 Diabetic Kidney Injury through PI3K/Akt Pathway. J Diabetes Res, 2023. 2023: p. 8810106.

44. Caetano-Pinto, P., et al., Cetuximab Prevents Methotrexate-Induced Cytotoxicity in Vitro through Epidermal Growth Factor Dependent Regulation of Renal Drug Transporters. Mol Pharm, 2017. 14(6): p. 2147–2157.

45. Ezzine, C., et al., Fatty acids produced by the gut microbiota dampen host inflammatory responses by modulating intestinal SUMOylation. Gut Microbes, 2022. 14(1): p. 2108280.

46. Wu, W., A.V. Dnyanmote, and S.K. Nigam, Remote communication through solute carriers and ATP binding cassette drug transporter pathways: an update on the remote sensing and signaling hypothesis. Mol Pharmacol, 2011. 79(5): p. 795–805.

47. Nigam, S.K. and K.T. Bush, Uraemic syndrome of chronic kidney disease: altered remote sensing and signalling. Nat Rev Nephrol, 2019. 15(5): p. 301–316.

48. Nieskens, T.T.G., et al., A Human Renal Proximal Tubule Cell Line with Stable Organic Anion Transporter 1 and 3 Expression Predictive for Antiviral-Induced Toxicity. The AAPS Journal, 2016. 18(2): p. 465–475.

49. Li, Y.J., et al., Short-chain fatty acids directly exert anti-inflammatory responses in podocytes and tubular epithelial cells exposed to high glucose. Front Cell Dev Biol, 2023. 11: p. 1182570.

50. Stein, R.A. and L. Riber, Epigenetic effects of short-chain fatty acids from the large intestine on host cells. microLife, 2023. 4.

51. Li, Y.J., et al., Dietary Fiber Protects against Diabetic Nephropathy through Short-Chain Fatty Acid-Mediated Activation of G Protein-Coupled Receptors GPR43 and GPR109A. J Am Soc Nephrol, 2020. 31(6): p. 1267–1281.

52. Kim, M.H., et al., Short-Chain Fatty Acids Activate GPR41 and GPR43 on Intestinal Epithelial Cells to Promote Inflammatory Responses in Mice. Gastroenterology, 2013. 145(2): p. 396–406.e10.

53. Kobayashi, M., et al., Short-chain fatty acids, GPR41 and GPR43 ligands, inhibit TNF-α-induced MCP-1 expression by modulating p38 and JNK signaling pathways in human renal cortical epithelial cells. Biochem Biophys Res Commun, 2017. 486(2): p. 499–505.

54. Martin-Gallausiaux, C., et al., Butyrate produced by gut commensal bacteria activates TGF-beta1 expression through the transcription factor SP1 in human intestinal epithelial cells. Sci Rep, 2018. 8(1): p. 9742.

55. Folkerts, J., et al., Butyrate inhibits human mast cell activation via epigenetic regulation of FcεRI-mediated signaling. Allergy, 2020. 75(8): p. 1966–1978.

56. Schilderink, R., et al., The SCFA butyrate stimulates the epithelial production of retinoic acid via inhibition of epithelial HDAC. Am J Physiol Gastrointest Liver Physiol, 2016. 310(11): p. G1138–46.

57. Gill, P.A., et al., Review article: short chain fatty acids as potential therapeutic agents in human gastrointestinal and inflammatory disorders. Aliment Pharmacol Ther, 2018. 48(1): p. 15–34.

58. Xie, H., et al., Uremic toxins mediate kidney diseases: the role of aryl hydrocarbon receptor. Cellular & Molecular Biology Letters, 2024. 29(1): p. 38.

59. Modoux, M., et al., Butyrate acts through HDAC inhibition to enhance aryl hydrocarbon receptor activation by gut microbiota-derived ligands. Gut Microbes, 2022. 14(1): p. 2105637.

60. Marumo, T., et al., Histone deacetylase modulates the proinflammatory and -fibrotic changes in tubulointerstitial injury. Am J Physiol Renal Physiol, 2010. 298(1): p. F133–41.

61. Liu, N., et al., Blocking the class I histone deacetylase ameliorates renal fibrosis and inhibits renal fibroblast activation via modulating TGF-beta and EGFR signaling. PLoS One, 2013. 8(1): p. e54001.

62. Pang, M., et al., Inhibition of histone deacetylase activity attenuates renal fibroblast activation and interstitial fibrosis in obstructive nephropathy. Am J Physiol Renal Physiol, 2009. 297(4): p. F996–f1005.

63. Hyndman, K.A., Histone Deacetylases in Kidney Physiology and Acute Kidney Injury. Semin Nephrol, 2020. 40(2): p. 138–147.

64. Shi, L., et al., HDAC6 Inhibition Alleviates Ischemia- and Cisplatin-Induced Acute Kidney Injury by Promoting Autophagy. Cells, 2022. 11(24).

65. Liu, J., et al., 2-methylquinazoline derivative F7 as a potent and selective HDAC6 inhibitor protected against rhabdomyolysis-induced acute kidney injury. PLoS One, 2019. 14(10): p. e0224158.

66. Tang, J., et al., Blockade of histone deacetylase 6 protects against cisplatin-induced acute kidney injury. Clin Sci (Lond), 2018. 132(3): p. 339–359.

67. Zhang, Y., et al., HDAC9-mediated epithelial cell cycle arrest in G2/M contributes to kidney fibrosis in male mice. Nat Commun, 2023. 14(1): p. 3007.

68. Liu, M., et al., Histone deacetylase 9 exacerbates podocyte injury in hyperhomocysteinemia through epigenetic repression of Klotho. Pharmacological Research, 2023. 198: p. 107009.

69. Lan, Z., et al., Downregulation of HDAC9 by the ketone metabolite β-hydroxybutyrate suppresses vascular calcification. J Pathol, 2022. 258(3): p. 213–226.

70. He, P., et al., Hdac9 inhibits medial artery calcification through down-regulation of Osterix. Vascul Pharmacol, 2020. 132: p. 106775.

71. Butler, P.L., A. Staruschenko, and P.M. Snyder, Acetylation stimulates the epithelial sodium channel by reducing its ubiquitination and degradation. J Biol Chem, 2015. 290(20): p. 12497–503.

72. Wang, Y., et al., Upregulation of histone acetylation reverses organic anion transporter 2 repression and enhances 5-fluorouracil sensitivity in hepatocellular carcinoma. Biochem Pharmacol, 2021. 188: p. 114546.

73. Vecsey, C.G., et al., Histone deacetylase inhibitors enhance memory and synaptic plasticity via CREB:CBP-dependent transcriptional activation. J Neurosci, 2007. 27(23): p. 6128–40.

74. Liu, W., et al., HDAC6 regulates epidermal growth factor receptor (EGFR) endocytic trafficking and degradation in renal epithelial cells. PLoS One, 2012. 7(11): p. e49418.

75. Mylroie, H., et al., PKCε-CREB-Nrf2 signalling induces HO-1 in the vascular endothelium and enhances resistance to inflammation and apoptosis. Cardiovascular Research, 2015. 106(3): p. 509–519.

76. Yu, L., et al., CREB1 protects against the renal injury in a rat model of kidney stone disease and calcium oxalate monohydrate crystals-induced injury in NRK-52E cells. Toxicol Appl Pharmacol, 2021. 413: p. 115394.

77. Yang, A., et al., Role of CREB1 dysregulation in calcium oxalate monohydrate crystals-induced tubular epithelial cell injury. Molecular & Cellular Toxicology, 2023.

78. Shan, Q., et al., Epigenetic modification of miR-10a regulates renal damage by targeting CREB1 in type 2 diabetes mellitus. Toxicol Appl Pharmacol, 2016. 306: p. 134–43.

79. Wang, D., et al., FOXO1 inhibition prevents renal ischemia-reperfusion injury via cAMP-response element binding protein/PPAR-γcoactivator-1α-mediated mitochondrial biogenesis. Br J Pharmacol, 2020. 177(2): p. 432–448.

80. Gao, L., et al., Taxifolin improves disorders of glucose metabolism and water-salt metabolism in kidney via PI3K/AKT signaling pathway in metabolic syndrome rats. Life Sci, 2020. 263: p. 118713.

81. Guan, T., et al., Troxerutin alleviates kidney injury in rats via PI3K/AKT pathway by enhancing MAP4 expression. Food Nutr Res, 2022. 66.

82. Wang, D., et al., FGF1(ΔHBS) ameliorates chronic kidney disease via PI3K/AKT mediated suppression of oxidative stress and inflammation. Cell Death Dis, 2019. 10(6): p. 464.

83. Kma, L. and T.J. Baruah, The interplay of ROS and the PI3K/Akt pathway in autophagy regulation. Biotechnol Appl Biochem, 2022. 69(1): p. 248–264.

84. He, J., et al., BCL2L10/BECN1 modulates hepatoma cells autophagy by regulating PI3K/AKT signaling pathway. Aging (Albany NY), 2019. 11(2): p. 350–370.

85. Lu, R., et al., Protective role of Astragaloside IV in chronic glomerulonephritis by activating autophagy through PI3K/AKT/AS160 pathway. Phytother Res, 2020. 34(12): p. 3236–3248.

86. Liu, P., et al., Crocetin attenuates the oxidative stress, inflammation and apoptosisin arsenic trioxide-induced nephrotoxic rats: Implication of PI3K/AKT pathway. International Immunopharmacology, 2020. 88: p. 106959.

