## Supplementary data for "Gut microbial-derived short chain fatty acids enhance kidney proximal tubule cell secretory function"

### Supplementary information

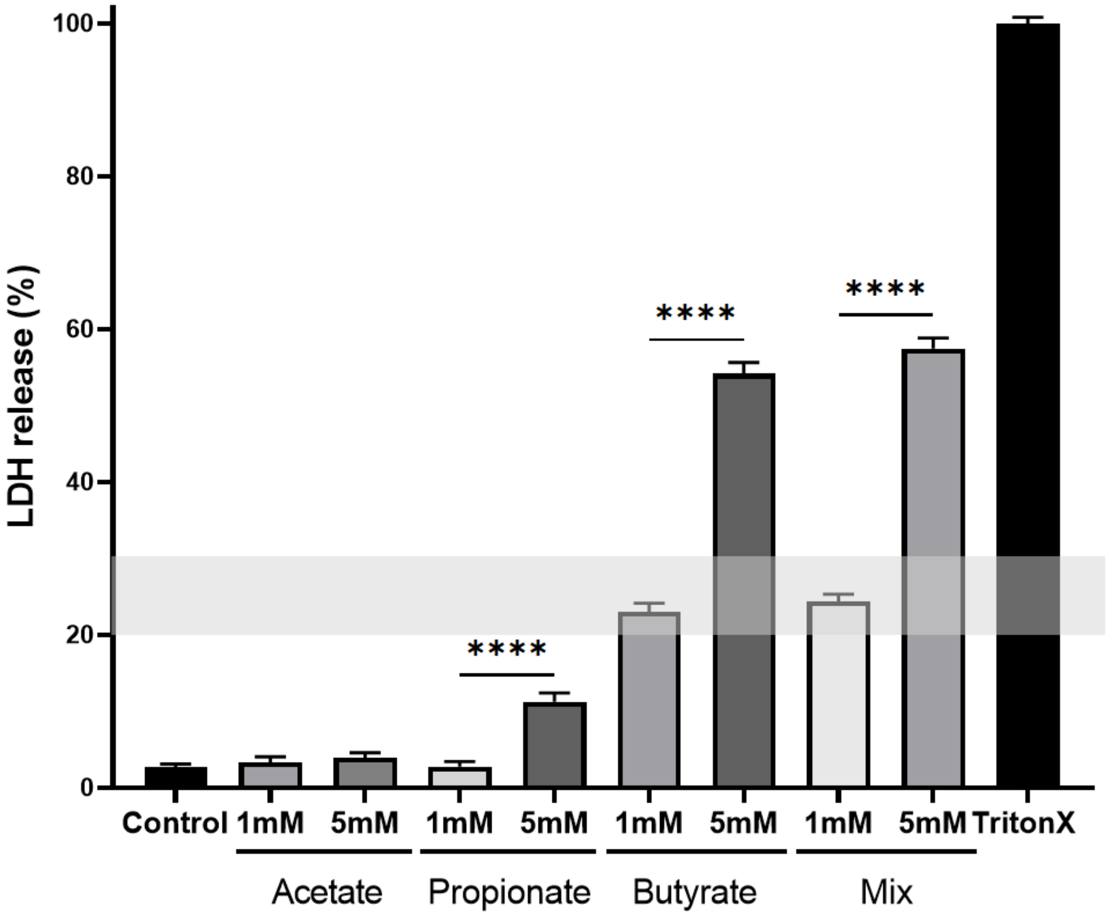


**Supplementary figure 1. SCFAs acetate, propionate and butyrate are relatively safe to ciPTECs-OAT1**. LDH cytotoxicity assay results displaying the % in LDH release when compared to control following treatment for 24 h with 1-5 mM of acetate, propionate and butyrate either alone or in combination. Results were calculated relatively to the positive control of 1% Triton inducing LDH release assigned as 100%. Values above the 20-30% window thresholds are considered toxic. All data is reported as mean ± SEM and were all statistically significant compared to the Triton X treatment ($p < 0.0001$). ****$p < 0{,}0001$ compared different concentration of the same treatment, as determined by Kruskal-Wallis test following Dunn's multiple comparison adjustment and Shapiro-Wilk test to assess for normality of data distribution.

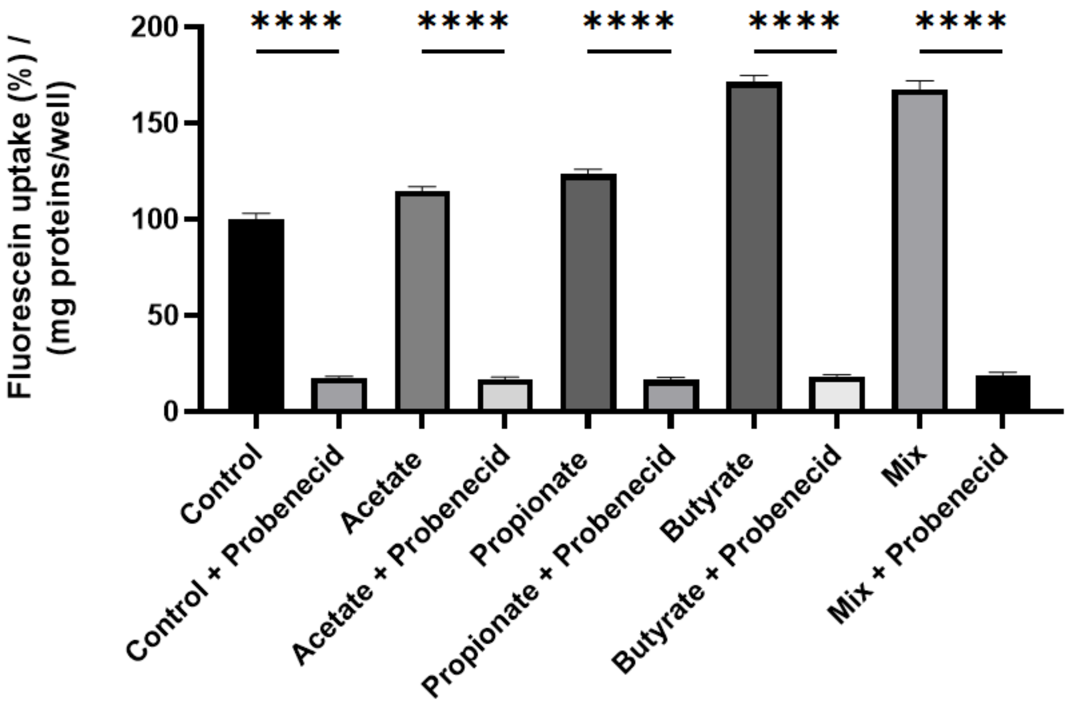


**Supplementary figure 2: Probenecid inhibit the SCFAs-mediated boosted OAT1 activity**. Activity of the OAT1 transporter in the presence of SCFAs alone or co-incubated with the OAT1 inhibitor Probenecid. No difference in OAT1 activity inhibition was observed upon exposure of ciPTEC-OAT1 to Probenecid with control or SCFAs. All data are representative of 4 independent experiments and are reported as mean ± SEM. Data were checked for normal distribution with Shapiro-Wilk test and statistical significance was analyzed with One-Way ANOVA followed by Posthoc Dunnet test (****$p$ < 0,0001).

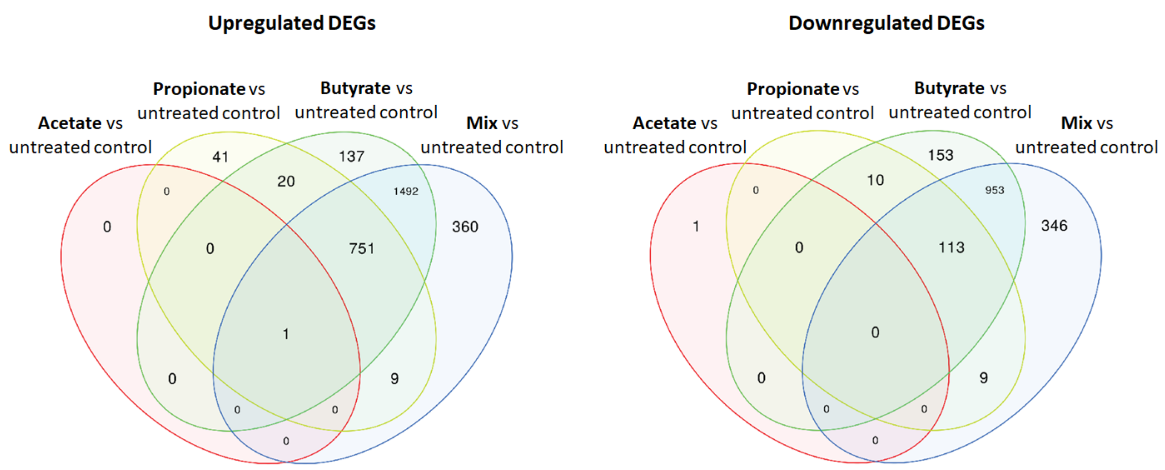


**Supplementary figure 3: Gene expression of upregulated and downregulated genes that are uniquely or shared between acetate, propionate, butyrate or mix treatments groups upon comparison to untreated control.** Venn diagram displaying the statistically significantly upregulated (left) and downregulated (right) DEGs (FC > 2, FDR < 0.05).

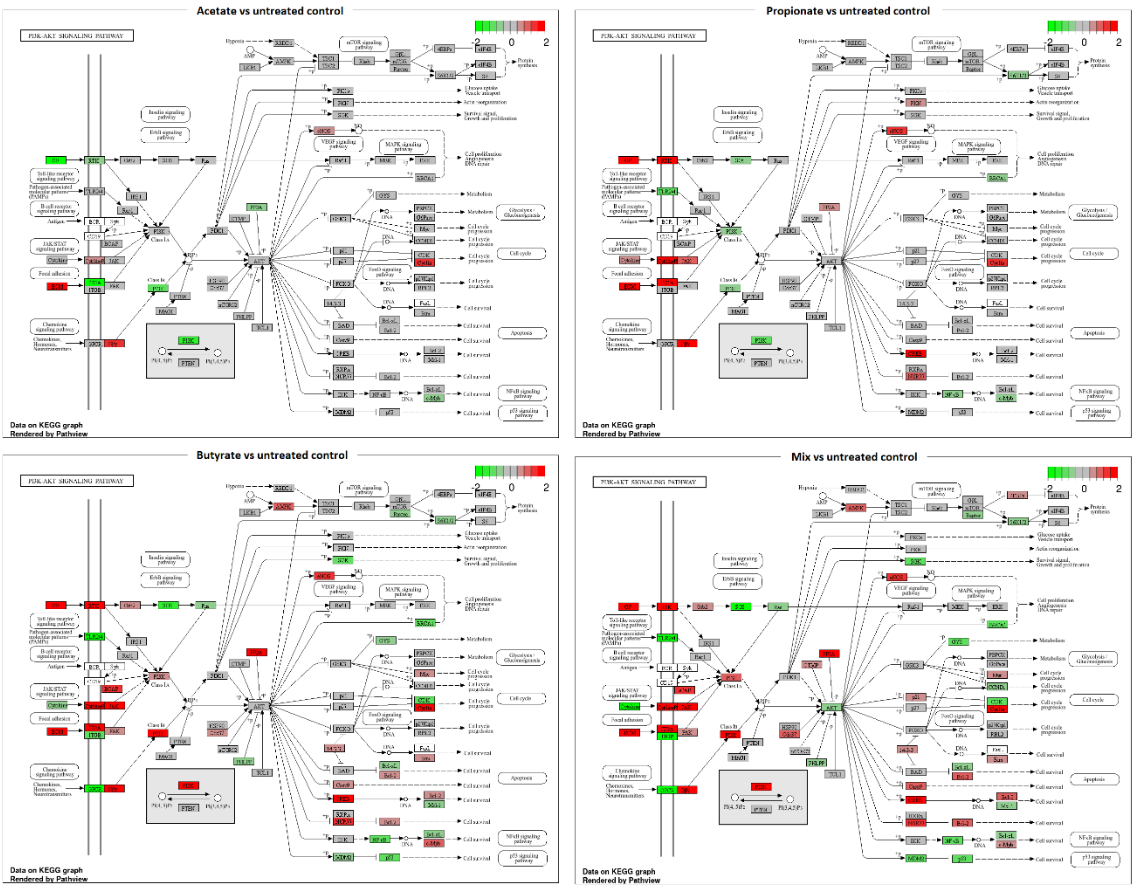


**Supplementary figure 4: SCFAs regulate PI3K-Akt signaling pathway markers.** Schematic representation of the PI3K-Akt signaling pathway upon exposure to acetate, propionate, butyrate or mix compared to the untreated control. Green: gene downregulation; red: gene upregulation.

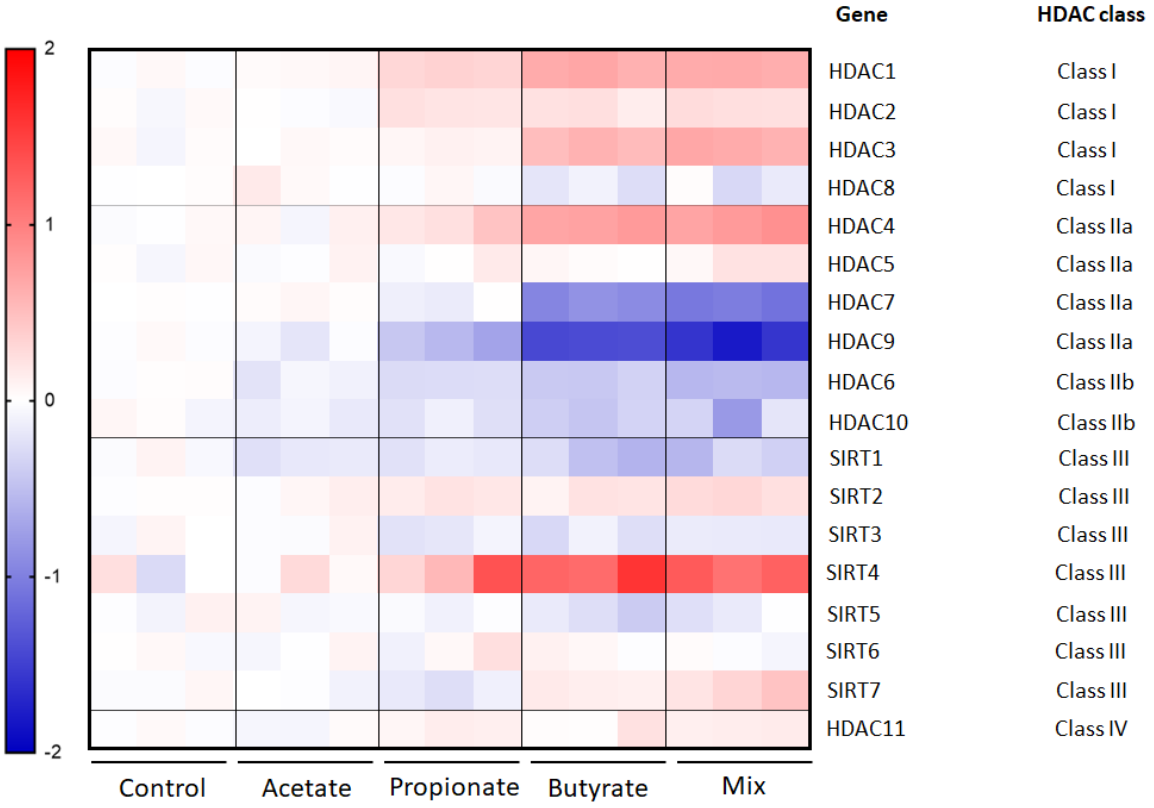


**Supplementary figure 5:** Heatmap displaying the 18 differentially expressed HDAC genes upon exposure of ciPTEC-OAT1 to acetate, propionate, butyrate and mix treatments compared to untreated control. Data is expressed as $Log_2$-Fold change.

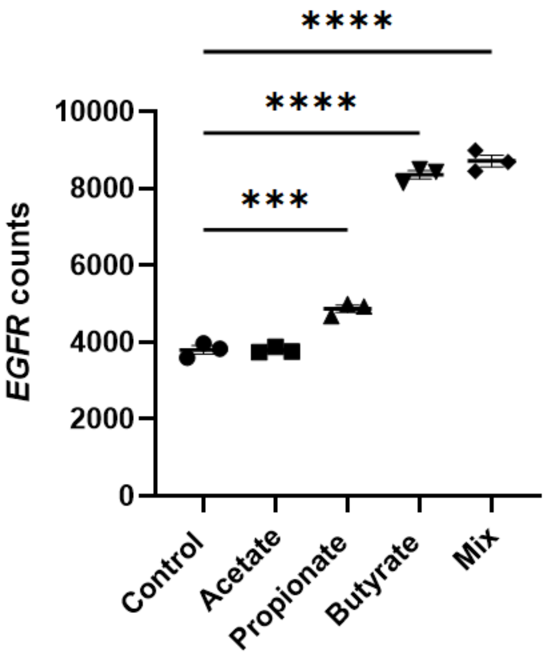


**Supplementary figure 6. EGFR gene counts**: Counts of the EGFR gene expression in the presence of acetate, propionate and butyrate either alone or in combination. Data is reported as mean ± SEM. ***$p$ < 0.001, ****$p$ < 0,0001 as determined by One-Way ANOVA following Dunnet's multiple comparison adjustment and Shapiro-Wilk test to assess for normality of data distribution.

**Supplementary information Westernblot**

OAT1 MEMBRANE

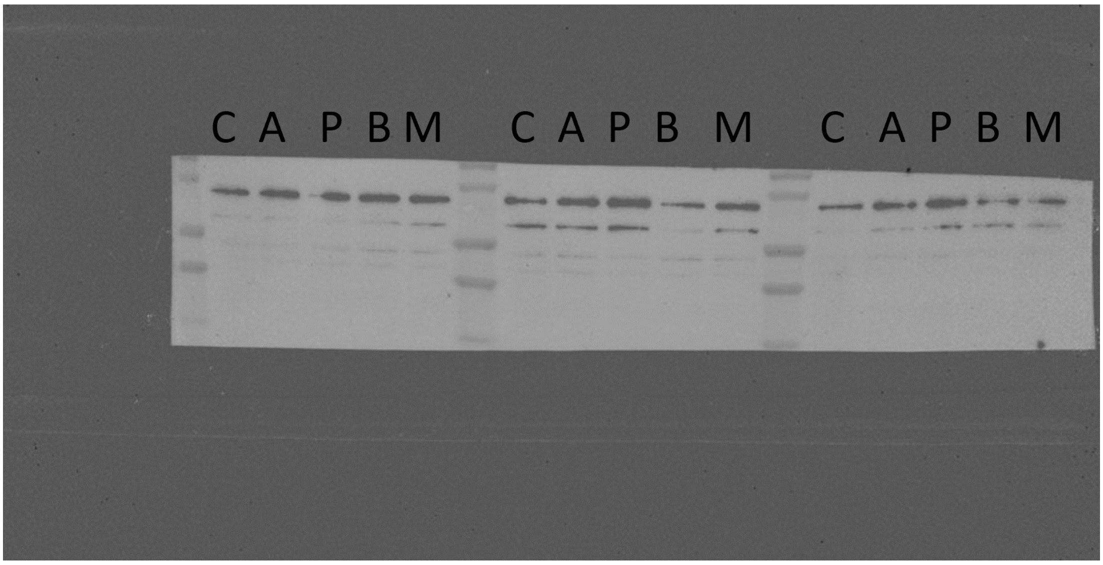


BETA TUBULIN MEMBRANE

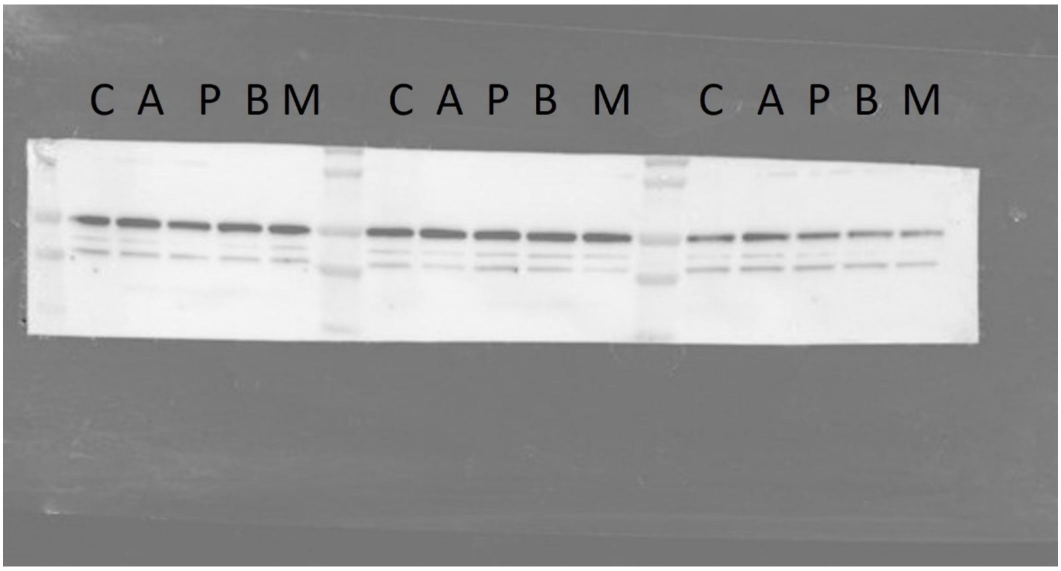


C = Control

A = Acetate

P = Propionate

B = Butyrate

M = Mix
